# Recapitulating idiopathic pulmonary fibrosis related alveolar epithelial dysfunction in an iPSC-derived air-liquid interface model

**DOI:** 10.1101/830109

**Authors:** Eva Schruf, Victoria Schroeder, Huy Q. Le, Tanja Schönberger, Dagmar Raedel, Emily L. Stewart, Katrin Fundel-Clemens, Teresa Bluhmki, Sabine Weigle, Michael Schuler, Matthew J. Thomas, Ralf Heilker, Megan J. Webster, Martin Dass, Manfred Frick, Birgit Stierstorfer, Karsten Quast, James P. Garnett

## Abstract

An abnormal emergence of airway epithelial-like cells within the alveolar compartments of the lung, herein termed bronchiolization, is a process often observed in patients suffering from idiopathic pulmonary fibrosis (IPF), a fatal disease characterized by progressive fibrotic lung remodeling. However, the origin of this dysfunctional epithelium remains unknown.

In this study, we aimed to investigate the effects of a pro-fibrotic milieu, similar to that found in an IPF lung, on human alveolar epithelial progenitor cell differentiation. We developed an induced pluripotent stem cell (iPSC)-derived air-liquid interface (ALI) model of alveolar type II (ATII)-like cell differentiation and stimulated it with an IPF-relevant cocktail (IPF-RC), composed of cytokines previously reported to be elevated in IPF lungs. iPSC-derived cultures express ATII markers and contain lamellar body-like structures. Stimulation with IPF-RC during the last two weeks of differentiation increases secretion of IPF biomarkers. Transcriptome analysis of IPF-RC treated cultures reveals significant overlap with human IPF data and enrichment of transcripts associated with extracellular matrix organization. IPF-RC stimulation further impairs ATII differentiation by driving a shift towards an airway epithelial-like expression signature.

In conclusion, we show for the first time, the establishment of a human model system that recapitulates aspects of IPF-associated bronchiolization *in vitro*. Our findings reveal how aberrant alveolar epithelial progenitor cell differentiation in a pro-fibrotic environment could contribute to alveolar bronchiolization in the distal IPF lung.

**SOURCE OF SUPPORT:** The research was funded by Boehringer Ingelheim Pharma GmbH & Co. KG.

## INTRODUCTION

Idiopathic pulmonary fibrosis (IPF) is a lethal respiratory disease characterized by progressive fibrosis of the lung parenchyma and lung function decline (1), and despite the emergence of new therapies in nintedanib and pirfenidone that can slow down disease progression, the disease remains inevitably fatal (2). Evidence suggests that age-related and genetic predisposition, as well as aberrant wound healing of lung epithelial micro-injuries and dysregulated fibro-proliferative repair are involved in IPF pathogenesis (3, 4). Dysfunctional epithelial repair in the IPF lung results in a loss of the normal proximo-distal patterning of the respiratory epithelium and an aberrant emergence of airway epithelial-like cell types within the alveolar regions of the lung, a phenomenon referred to as bronchiolization (5, 6). Intermediate cells types, co-expressing alveolar and conducting airway cell selective markers, reside in these bronchiolized regions, indicative of aberrant lung epithelial differentiation (7). Yet the definite origin of the bronchiolized epithelial cells lining the microscopic honeycomb cysts of IPF patients remains unknown.

The identification of specific bronchoalveolar cells as putative stem cells of the distal lung in mouse models of lung injury together with the observation of increased numbers of basal cell and club cell populations in IPF lesions have led to the hypothesis that the bronchiolized epithelium in IPF might originate from airway basal or bronchoalveolar cell migration towards alveolar regions (8-11). However, the role of bronchoalveolar stem cells in human lung homeostasis and repair remains debatable and several recent reports strongly suggest that a specified Wnt responsive alveolar epithelial stem cell population, constituting a subpopulation of alveolar type II (ATII) cells residing within a specific mesenchymal niche, serves as the residual stem cell pool of the alveolar epithelium (12-16). In line with this, it has been hypothesized that the susceptibility of surfactant secreting ATII cells to injury and the resulting defective alveolar repair could play a major role in IPF disease onset and progression (15, 17, 18).

Limited access to primary human ATII cells, particularly from diseased patients, and the rapid loss of ATII marker expression and function when cultured *in vitro*, make it challenging to study the role of human ATII cells in IPF (19). Recent advances have led to the successful derivation of ATII-like cells from human pluripotent stem cells (hPSCs, including both embryonic stem cells and induced pluripotent stem cells), providing a promising alternative cell source for *in vitro* disease modeling (20-26). Previous reports on hPSC-derived lung cells showed that despite their phenotypic similarities to mature lung epithelium, these models are equivalent to human fetal lung cells, rather than representing an adult state (27, 28). We propose that hPSC-derived lung epithelial cultures constitute a highly valuable model to gain insights into the role of stem cells during lung repair and regeneration due to their immature nature by which they resemble adult alveolar epithelial progenitors (AEPs), a Wnt responsive subpopulation of ATII cells recently identified as the alveolar stem cell in the human lung that shows enrichment of genes associated with lung development (16). This is in line with previous reports, highlighting the influence of developmental pathways in lung regeneration (32-34). Moreover, a role for aberrantly activated developmental pathways, including Wnt, SHH, Notch and FGF signaling, has also been described in the pathogenesis of IPF, the progression of which is commonly viewed as a vicious cycle of lung injury and repair (35-37).

In this study, we sought to test the hypothesis that alveolar epithelial stem cells exposed to a pro-fibrotic environment could contribute to the metaplastic airway-like epithelium lining IPF cystic lesions. Building on previous efforts to derive alveolar epithelium from hPSCs, we aimed to develop a novel model system that allows human ATII-like differentiation from induced pluripotent stem cells (iPSCs) in 2D air-liquid interface (ALI) culture. We next aimed to apply this model to studying alveolar epithelial dysfunction in IPF by exposing iPSC-derived alveolar epithelial progenitor cells to an IPF-relevant cocktail (IPF-RC), based on upregulated cytokines found in IPF patient bronchoalveolar lavage or sputum (38-46).

## METHODS

Human iPSC lines SFC065-03-03 (EBiSC: STBCi057-A, Biosamples ID: SAMEA104493762) and SFC084-03-01 (EBiSC: STBCi033-A, Biosamples ID: SAMEA104493681) were obtained from the StemBANCC consortium. Primary human Small Airway Epithelial Cells (CC-2547, Lot 501937) and primary human lung fibroblasts (CC-2512, Lot. 0000608197) were obtained from Lonza (Basel, Switzerland).

Formalin-fixed and paraffin embedded lung samples from human IPF patients were purchased from Folio Biosciences (Powell, OH, US) under the regulatory conditions of the Boehringer Ingelheim corporate policy regarding the acquisition and use of human biospecimen. Samples were reviewed internally by a trained pathologist and the initial diagnosis of IPF was confirmed.

We developed a directed differentiation protocol to derive ATII-like cells from human iPSCs in air-liquid interface (ALI) culture. A detailed description of the differentiation protocol is provided in **SI Appendix, SI Materials and Methods**.

A novel IPF-relevant cocktail (IPF-RC) was designed to model aspects of the IPF-related lung milieu *in vitro*. A panel of nine cytokines, previously reported to be upregulated in clinical IPF BAL or sputum samples compared to healthy control lungs, was selected based on a literature research. The composition of IPF-RC is summarized in **SI Appendix, Table E1**.

All antibodies are listed in **SI Appendix, Table E2** and TaqMan Gene Expression Assays are listed in **Table E3**. RNA-seq data is available in **SI Supplementary RNA-seq data**.

Extended experimental procedures are described in **SI Appendix, SI Materials and Methods**.

## RESULTS

### Differentiation of human iPSCs towards ATII-like cells in ALI culture

We established a directed step-wise differentiation protocol to derive ATII-like cells from human iPSCs, in which distal lung progenitor maturation towards ATII-like cells was carried out on transwell inserts at air-liquid interface (ALI) to mimic the physiological environment of the mature alveolar epithelium (**Figure 1**A). Adapting a previously published protocol (20, 47), we generated NKX2.1^+^/ FOXA2^+^/ SOX9^+^ lung epithelial progenitor cells, via SOX17^+^ definitive endoderm and ventralised anterior foregut endoderm (**Figure 5B, C, D; Figure S1A, B**). 3D matrigel culture was utilized to confirm the budding and branching potential of the iPSC-derived lung epithelial progenitor cells, which constitutes a key functional feature of fetal epithelial progenitors during lung morphogenesis (**Figure S1C**). Subsequently, to induce ATII-like differentiation, day 24 lung epithelial progenitor cells were seeded onto transwell inserts and confluent cultures were further differentiated at air-liquid-interface until day 49, with temporal modulation of Wnt signaling applied to drive further differentiation of NKX2.1^+^ lung progenitor cells towards ATII-like cells (23). *SFTPC, SFTPB* and *ABCA3* expression was induced over time (**Figure 5E**), together with an upregulation of many other ATII associated transcripts by day 49 (**Figure 5F**). Cells differentiated at air-liquid-interface displayed upregulated expression levels of *SFTPC, SFTPB* and *ABCA3* compared to submerged conditions, while no induction of the ATI markers *CAV1* or *PDPN* was observed under either condition (**Figure 6A**). Immunofluorescence showed the homogeneous expression of the pan-epithelial marker E-Cadherin and confirmed the presence of SFTPC^+^, SFTPB^+^ and ABCA3^+^ cells, indicative of an ATII cell phenotype (**Figure 6B**). Day 49 cultures also showed upregulated expression of the human alveolar epithelial progenitor marker *TM4SF1* compared to iPSCs. Moreover, a moderate induction of basal cell markers (*KRT5, TP63*) and other airway related transcripts (*SCGB1A1, MUC5B* and *FOXJ1*) was observed and rare individual KRT5^+^ cells could be detected, indicating that in addition to ATII-like cells, our model system contains airway basal-like cells (**Figure S2A, B**). Staining of cross-sectioned day 49 transwell cultures revealed that SFTPC^+^ cells resided mainly on the apical air-exposed surface of the cultures (**Figure 6C**). ATII-specific lamellar body-like structures were observed (**Figure 6D**) and live-stained iPSC-derived ATII-like cells contained large organelles displaying intensive accumulation of LysoTracker Green DND-26 (**Figure 6E**). Single cell dissociation and replating of iPSC-derived ATII-like cells under 2D submerged conditions in serum-containing medium resulted in downregulated *SFTPC, SFTPB* and *ABCA3* expression and upregulated expression of the ATI markers *CAV1* and *PDPN*, consistent with ATI-like differentiation potential (**Figure S2C**). Our ATII differentiation protocol was reproducible in an alternative human iPSC line (**Figure S3**).

**Figure 1.**
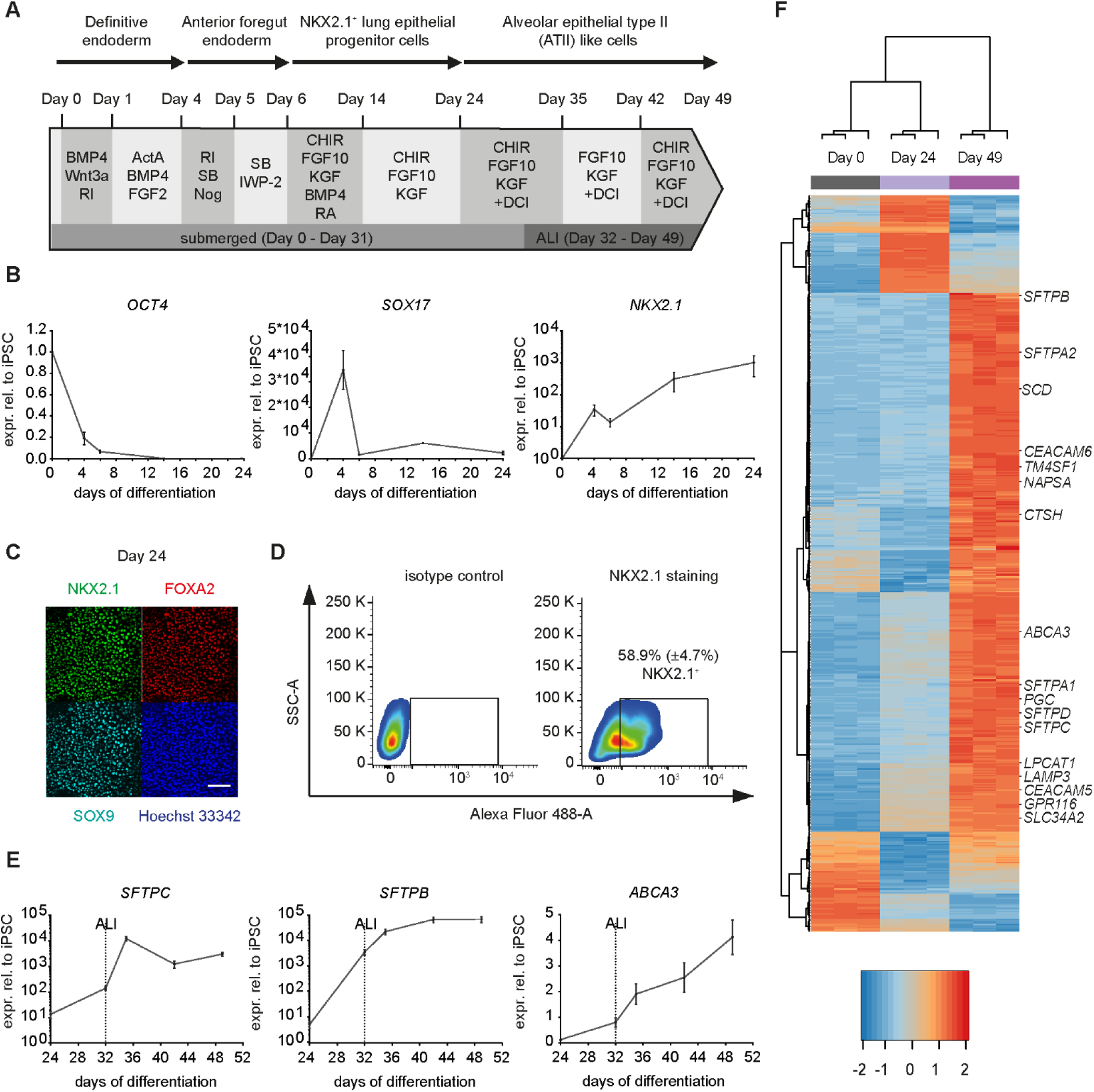
Transcriptional changes characteristic of distal lung development induced during directed differentiation of human iPSCs. **(A)** iPSC (line SFC065-03-03) to ATII-like cell differentiation protocol. RI= Y-27632; ActA = Activin A; SB = SB431542; Nog = Noggin; RA = all-trans retinoic acid; CHIR = CHIR99021; DCI = dexamethasone, 8-Br-cAMP, IBMX. (**B)** Transcription factor gene expression from day 0 to day 24 of differentiation normalized to GAPDH relative to day 0. Mean fold change ± SEM. N=3. **(C)** Immunofluorescence of day 24 lung progenitors stained against NKX2.1, FOXA2 and SOX9. Nuclei stained with Hoechst 33342. Scale bar 50µm. **(D)** Flow cytometry analysis of NKX2.1^+^ cells on day 24. Mean percentage of NKX2.1^+^ cells (± SEM) is indicated. N=3. **(E)** ATII marker gene expression from day 24 to day 49 of differentiation normalized to GAPDH relative to day 0. ALI = start of air-liquid interface. Mean fold change ± SEM. N=3. **(F)** Heatmap of top deregulated genes in day 49 vs. day 24 cultures by RNA-seq (cut-off: q-val < 0.001, |logFC| > 2, maxRPKM > 10) with known ATII marker genes labelled on the right. Day 0 = iPSCs. Day 24 = lung epithelial progenitors. Day 49 = ATII-like cells.

**Figure 2.**
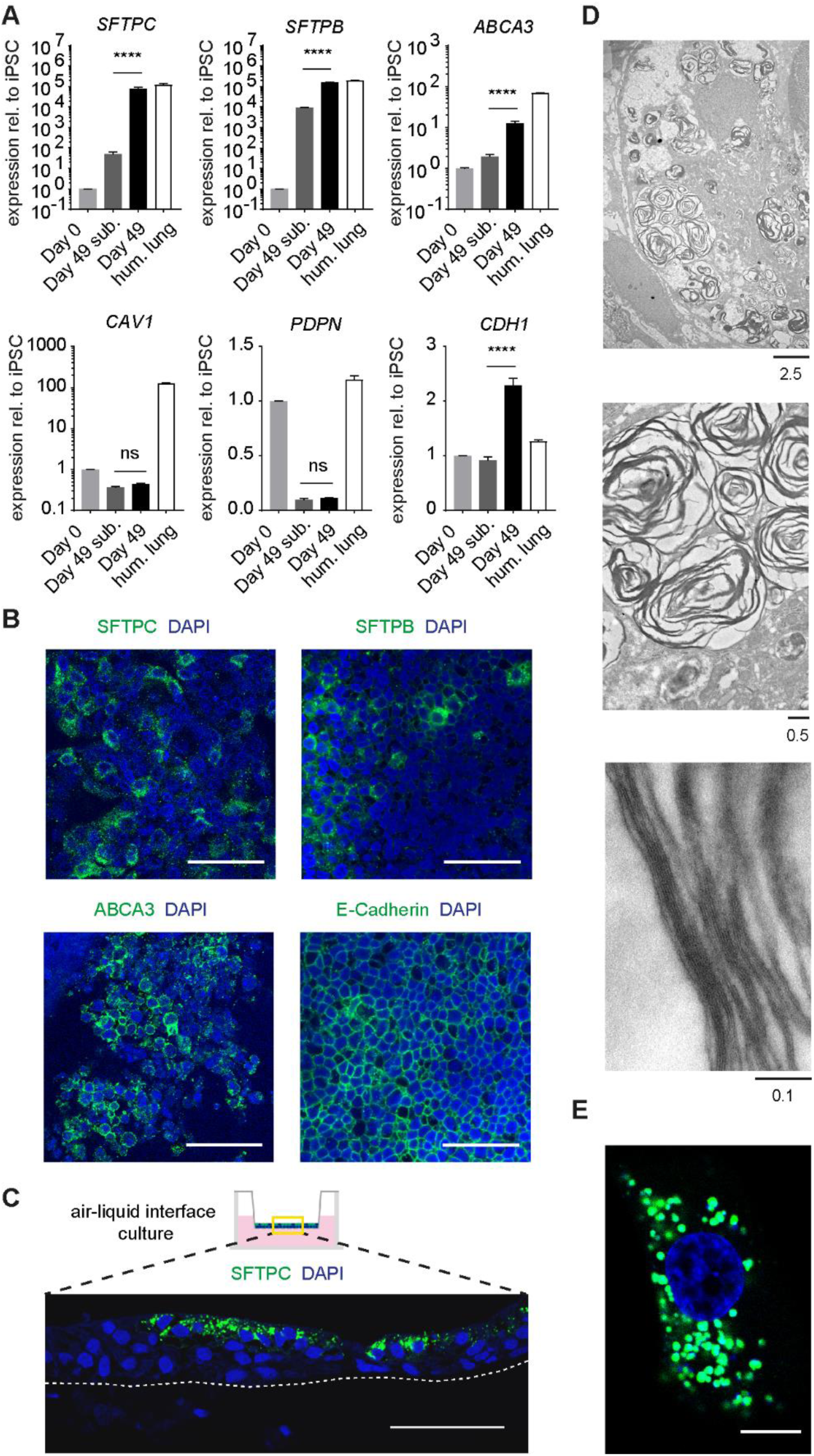
ATII-like phenotype of iPSC-derived air-liquid interface (ALI) cultures. **(A)** Alveolar epithelial marker gene expression on day 49 of differentiation normalized to GAPDH relative to day 0. Day 49 sub. = submerged conditions. Day 49 = ALI from day 32 of differentiation onwards. Hum. lung = total human lung control. Mean fold change ± SEM. N=3. ns not significant,**** P < 0.0001 by Mann-Whitney U-test. **(B)** Immunofluorescence of day 49 cultures stained against SFTPC, SFTPB, ABCA3 or E-Cadherin. Nuclei stained with DAPI. Scale bars 50µm. **(C)** Immunofluorescence of a day 49 ALI culture cross-section stained against SFTPC. Nuclei stained with DAPI. Dashed line indicates PET membrane of transwell insert. Scale bar 50 µm. **(D)** Lamellar body-like structures in day 49 cultures identified by transmission electron microscopy. Scale bars in µm. **(E)** LysoTracker Green live-staining of iPSC-derived ATII-like cell. Nucleus stained with Hoechst 33342. Scale bar 10 µm.

**Figure 3.**
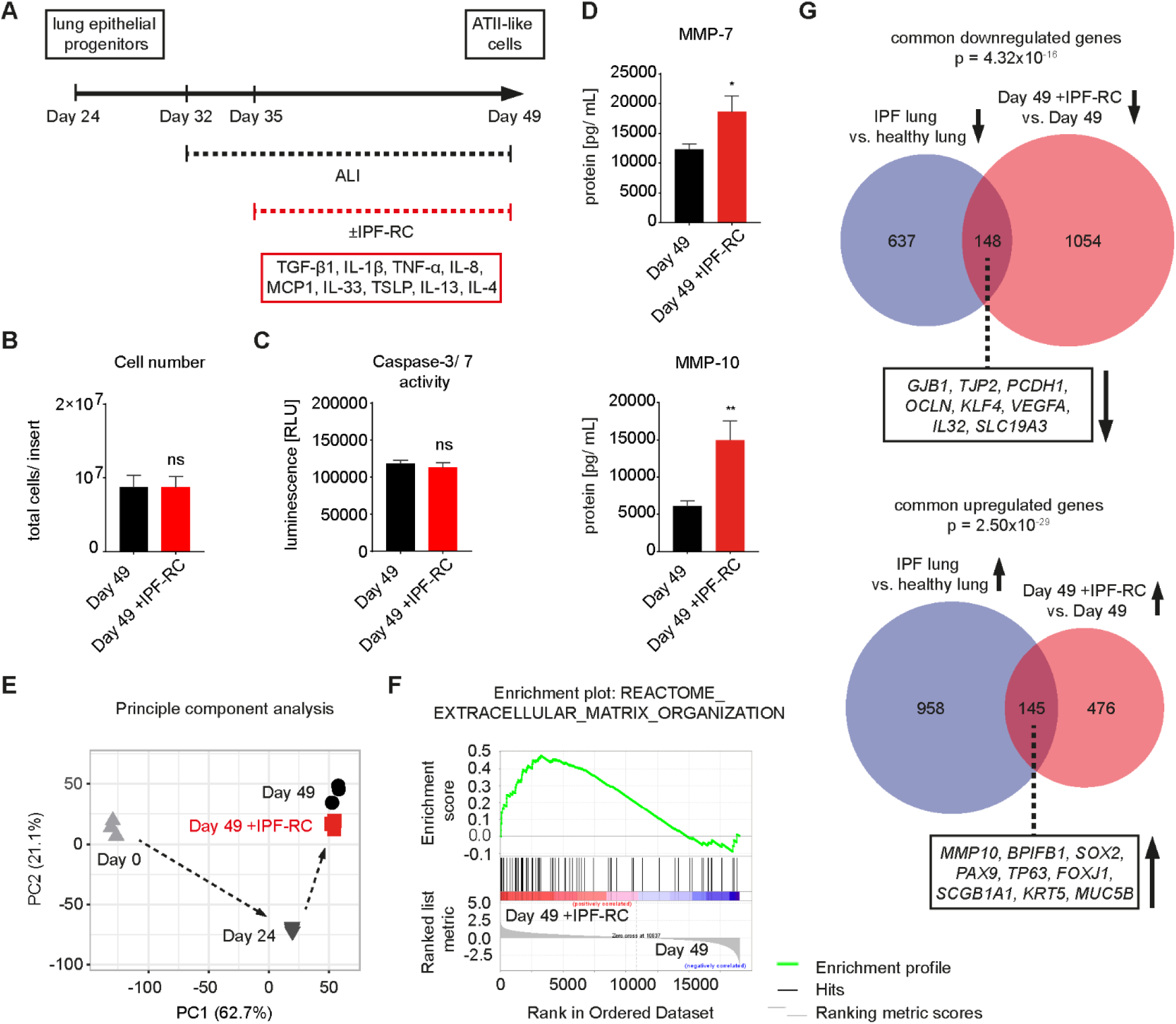
Fibrosis-related changes in transcriptional and secretory state caused by differentiation of iPSC-derived lung epithelial progenitors in the presence of an IPF-relevant cytokine cocktail (IPF-RC). **(A)** IPF-RC stimulation during alveolar epithelial differentiation (day 35 - day 49). **(B)** Total cell number per transwell insert. Day 49 = iPSC-derived ATII-like cells (control). Day 49 +IPF-RC = iPSC-derived cells treated with IPF-RC from day 35 of differentiation onwards. Mean ± SEM. N=4. ns not significant by Mann-Whitney U-test. **(C)** Caspase-3/ 7 activity. Mean ± SEM. N=3. ns not significant by Mann-Whitney U-test. **(D)** MMP-7 and MMP-10 protein levels in culture medium measured by ELISA. Mean ± SEM. N=3. * P < 0.05, ** P < 0.01 by Mann-Whitney U-test. **(E)** Principal component analysis of RNA-seq data representing the top 5000 transcripts according to the coefficient of variation. Arrows represent experimental timecourse. **(F)** Enrichment plot for the extracellular matrix organization gene set from Reactome. FDR q-value = 0.039. **(G)** Intersection between deregulated transcripts in IPF-RC treated iPSC-derived day 49 cultures (vs. untreated) and deregulated transcripts in human IPF lung tissue (vs. healthy). Cut-off: q-val < 0.05, |logFC| > 0.5, maxRPKM > 2. Top section represents down-regulated, bottom section up-regulated transcripts. Boxes contain examples of commonly up- or down-regulated transcripts.

**Figure 4.**
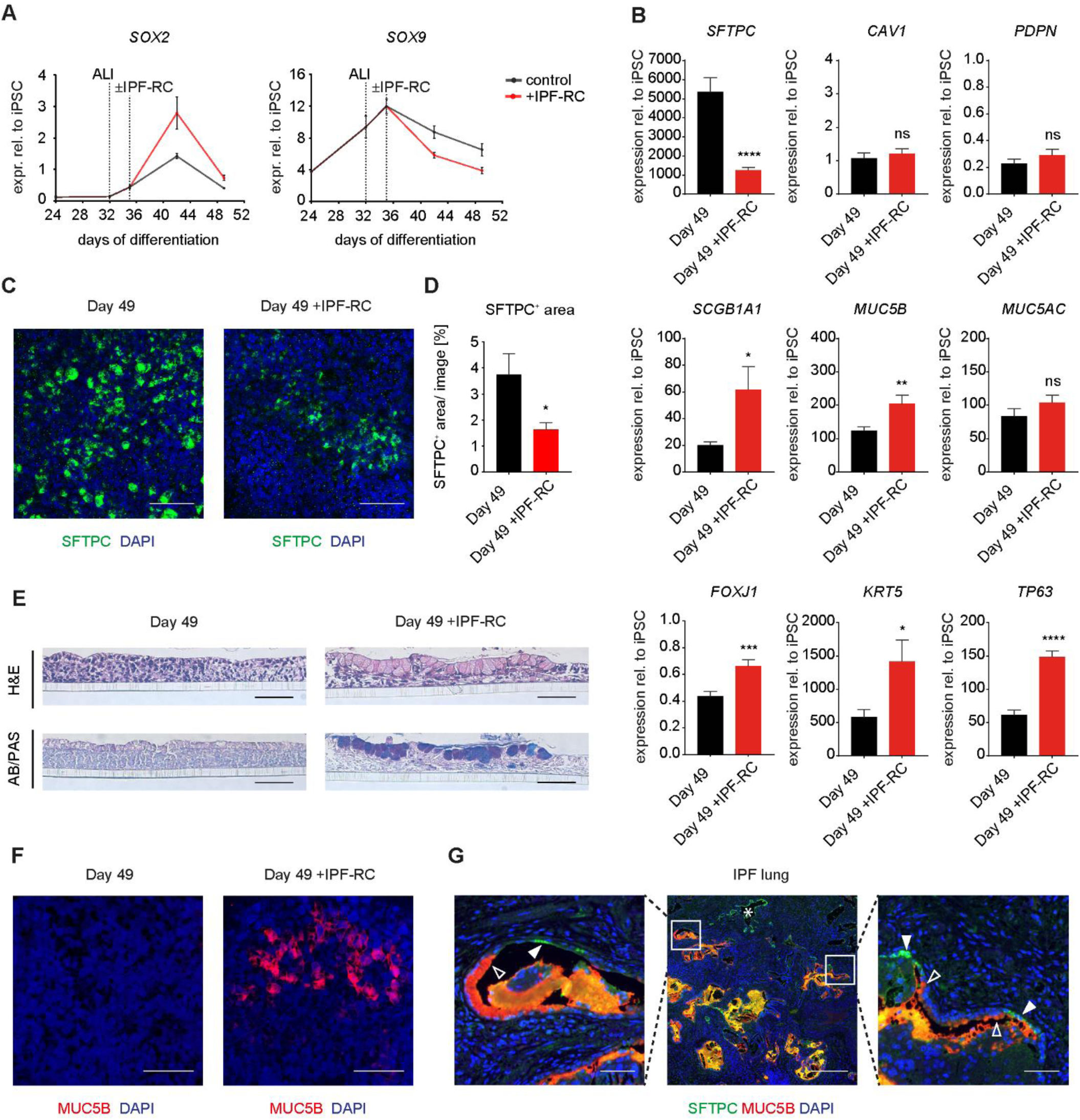
IPF-RC induced distal-to-proximal shift of iPSC-derived lung epithelial progenitor cell differentiation resulting in an epithelial phenotype mimicking aspects of IPF-related bronchiolization. **(A)** SOX2 and SOX9 expression from day 24 to day 49 of differentiation normalized to GAPDH relative to day 0. ALI = start of air-liquid interface. ±IPF-RC = start of IPF-RC/vehicle treatment. Mean fold change ± SEM. N=3. **(B)** Alveolar and airway epithelial marker expression in day 49 cultures ±IPF-RC treatment normalized to GAPDH relative to day 0. Mean fold change ± SEM. N=9. ns not significant, * P < 0.05, ** P < 0.01, *** P < 0.001 and **** P < 0.0001 by two-tailed Student’s t-test. **(C)** Immunofluorescence against SFTPC in day 49 cultures ±IPF-RC. Nuclei stained with DAPI. Scale bars 50µm. **(D)** SFTPC^+^ surface area quantified by immunofluorescence. Mean SFTPC^+^ area per image ± SEM. N=3. (n=3 biological replicates per differentiation round and 9 images per replicate). * P < 0.05 by Mann-Whitney U-test. **(E)** H&E and Alcian Blue/PAS stain of iPSC-derived culture cross-sections. Scale bars 50 µm. **(F)** Immunofluorescence against MUC5B in day 49 cultures ±IPF-RC. Nuclei stained with DAPI. Scale bars 40µm. **(G)** Immunofluorescence of human IPF lung against SFTPC and MUC5B. Nuclei stained with DAPI. Center: Cystic lesions in the distal lung. * marks lesion lined by simple SFTPC^+^ epithelium. Boxes mark lesions lined by metaplastic transitional epithelium. Scale bar 350µm. Left and right: Transition zones containing SFTPC^+^ cells (filled arrow heads) in close proximity to MUC5B^+^ cells (unfilled arrow heads). Scale bars 50µm.

### iPSC-derived distal lung epithelial progenitor differentiation in the presence of an IPF-relevant cytokine cocktail (IPF-RC) results in IPF-related changes in expression signature and secretory state

We utilized our iPSC-derived model system to investigate the effects of a pro-fibrotic environment on ATII cell differentiation. An IPF-relevant cytokine cocktail (IPF-RC) was designed and was capable of inducing collagen I formation in primary human lung fibroblasts (**Figure S4**). The IPF-RC, contains nine cytokines previously shown to be upregulated in IPF bronchoalveolar lavage or sputum samples compared to healthy controls (TGF-β1, IL-1β, TNF-α, IL-8, MCP1, IL-33, TSLP, IL-13 and IL-4; **Table S1**). IPF-RC was applied to iPSC-derived alveolar epithelial progenitor cells for two weeks, following the induction of *SFTPC* expression (**Figure 5E; Figure 7A**). No changes in total cell number or caspase-3/7 activity were detected upon IPF-RC treatment, indicating that IPF-RC did not exert any anti-proliferative or pro-apoptotic effects (**Figure 7B, C**). qRT-PCR revealed an upregulated expression of the IPF-related marker genes *MMP-7, MMP-10* and *BPIFB1* and significantly increased MMP-7 and MMP-10 protein levels were detected in the culture medium of day 49 cultures stimulated with IPF-RC by ELISA, while the expression of *CDH1, ACTA2* and *COL1A1* was not altered (**Figure 7D; Figure S5**).

RNA-seq was performed to further investigate IPF-RC mediated transcriptional changes (**Figure 7E**). In accordance with an upregulation of MMP-7 and MMP-10 secretion, gene set enrichment analysis revealed a significant enrichment of transcripts associated with extracellular matrix organization (among top 10 associated gene sets under investigation; C2.CP.REACTOME (Curated)) for the day 49 +IPF-RC phenotype (**Figure 7F**). We also compared the expression data from our iPSC-derived model to a publicly available dataset for human IPF and healthy lung tissue (48). Calculation of the intersection of these data sets revealed a significant overlap of both downregulated (p = 4.32×10^−16^) and upregulated (p = 2.5 × 10^−29^) transcripts between day 49 cultures treated with IPF-RC and human IPF lungs (**Figure 7G**). Epithelial cell adhesion related transcripts (*GJB1, TJP2, PCDH1* and *OCLN*), as well as transcripts that are known to be repressed in IPF, including *KLF4, VEGFA, IL32* and *SLC19A3*, were among the downregulated genes, while MMP10 and the airway epithelial associated transcripts *BPIFB1, SOX2, PAX9, TP63, FOXJ1, SCGB1A1* and *KRT5*, were upregulated in both datasets. These results indicate that IPF-RC stimulation of our iPSC-derived ALI model system allows for the recapitulation of IPF-relevant processes and transcriptional changes, *in vitro*.

### IPF-RC stimulation induces a shift in iPSC-derived distal lung epithelial progenitor cell differentiation towards a proximalized phenotype

As RNA-seq revealed that known airway epithelial related transcripts were among the upregulated transcripts upon IPF-RC stimulation, we sought to determine if treatment with IPF-RC had the potential to alter the proximo-distal differentiation pattern of iPSC-derived lung epithelial progenitors. Elevated expression of the transcription factor *SOX2* and reduced expression of *SOX9* were observed upon IPF-RC stimulation, both of which are central players in proximo-distal epithelial patterning during lung development (**Figure 8A**). IPF-RC treated cultures displayed significantly reduced expression of ATII-specific *SFTPC*, while the airway related transcripts *SCGB1A1, MUC5B, FOXJ1, KRT5* and *TP63*, which are also expressed in the abnormal epithelium lining cystic lesions found in distal IPF lungs, were significantly upregulated (**Figure 8B**). *MUC5AC* remained unchanged upon IPF-RC treatment, indicating that goblet cell associated mucin induction was specific to MUC5B, similar to the bronchiolized epithelium in IPF patients. Expression of *CAV1* and *PDPN* was not altered, indicating that differentiation of ATII-like cells towards ATI-like cells did not contribute to the loss of *SFTPC* expression.

To understand the relative contribution of mediators within the IPF-RC that are known to induce epithelial remodeling, individual components were removed from the cocktail. Day 49 cultures differentiated in the presence of IPF-RC without IL-13 expressed higher levels of *SFTPC* than cells treated with the full cytokine cocktail, and displayed lower expression of *MMP-10, BPIFB1, MUC5B, FOXJ1* and *TP63*, suggesting IL-13 as the predominant driver of the IPF-related effects exerted by IPF-RC (**Figure S6A**). However, removal of IL-13 did not reduce the expression of *MMP-7, SCGB1A1* and *KRT5* compared to the full cocktail. Furthermore, stimulation with 2.5 ng/mL IL-13 alone resulted in a significant downregulation of *SFTPC* expression and upregulation of *FOXJ1* and *TP63* expression, but did not alter the expression of *MMP-10, BPIFB1* or *MUC5B*, indicating that IL-13 was not solely responsible for the proximalization effect of IPF-RC (**Figure S6B**). IPF-RC without TGF-β1 led to a reduction in MUC5B expression compared to the full cocktail, yet failed to reduce IPF-RC mediated expressional changes in any of the other transcripts analyzed (**Figure S6A**). Removal of TNF-α did not result in a reduced induction of any IPF- or airway-associated transcripts compared to IPF-RC (**Figure S6A**). Moreover, TGF-β1 or TNF-α removal failed to rescue *SFTPC* expression (**Figure S6A**). These results indicate that TGF-β1 and TNF-α were not the main drivers of the bronchiolization-like effect cause by IPF-RC in our model system.

Nintedanib, a tyrosine kinase inhibitor known to slow down IPF-related lung function decline, only showed some minor effects on *SCGB1A1* and *TP63* expression at clinically relevant concentrations (100 nM), but was neither able to rescue induction of IPF- and other airway-related marker genes, nor the reduction in *SFTPC* expression in IPF-RC treated cells (**Figure S7**).

In accordance with the reduced *SFTPC* expression in IPF-RC cultures, the mean surface area covered by SFTPC^+^ cells was significantly reduced in day 49 IPF-RC cultures compared to controls, indicating a loss of ATII-like cells (**Figure 8C, D**). Histological analysis of cross-sectioned day 49 cultures revealed that IPF-RC stimulation resulted in the emergence of Alcian Blue/PAS^+^ cell clusters, displaying a secretory cell-like morphology (**Figure 8E**). Moreover, areas of MUC5B^+^ goblet-like cells, which were absent in controls, were detected in IPF-RC cultures by immunofluorescence (**Figure 8F**). These results show that IPF-RC led to the emergence of MUC5B^+^ secretory cells, which are also frequently observed lining cystic lesions in the distal lung of IPF patients, at the expense of ATII-like cell differentiation.

### IPF-RC treatment during primary small airway basal cell differentiation favors MUC5AC^+^ goblet cell and impairs ciliary cell formation

As airway basal cells have been proposed as a source of the bronchiolized distal lung epithelium in IPF, we also assessed the effect of IPF-RC treatment on primary small airway basal cell differentiation at air-liquid interface. Similar to our iPSC-derived cultures, IPF-RC stimulation induced upregulated expression of *KRT5, TP63, MMP7, MMP10* and *BPIFB1*, as well as increased secretion of MMP-7 and MMP-10 in small airway epithelial cell (SAEC) cultures (**Figure S8A-C**). Conversely, expression of *SCGB1A1* and *FOXJ1* decreased compared to control cultures and *MUC5AC* expression was enhanced, while no significant deregulation of *MUC5B* expression could be detected, the upregulation of which has been linked to epithelial dysfunction in IPF. Cilia formation and ciliary function were also severely impaired in IPF-RC treated small airway epithelium (**Figure S8D**). These results indicate that the effect of IPF-RC on primary small airway basal cell differentiation differs from its effect on iPSC-derived alveolar epithelial progenitor cell differentiation.

### The metaplastic epithelium lining cystic lesions in IPF lungs contains atypical transition zones between SFTPC^+^ ATII-like cells and MUC5B^+^ goblet-like cells

To confirm the relevance of the findings observed in our model system for human disease, immunofluorescence in IPF patient lung sections was performed. The heterogeneous cystic lesions found in the dense fibrotic tissue of the distal IPF lungs were lined by a simple single-layered columnar or cuboidal epithelium, interrupted by areas of epithelial denudation and areas of multi-layered hyperplastic epithelium. Double immunofluorescence against SFTPC and MUC5B was performed in three IPF lungs. In accordance with previous reports that identified MUC5B^+^ secretory cells as the predominant mucus cell type in the honeycomb epithelium, many of the cystic lesions found in the distal regions of IPF lungs were lined by MUC5B expressing columnar epithelial cells and displayed extensive mucus plugging. Immunofluorescence further confirmed the presence of SFTPC^+^ cells in distal IPF patients’ lungs, with some cystic lesions lined exclusively by a simple SFTPC^+^ epithelial cell layer, indicative of ATII cell hyperplasia (**Figure 8G)**. The luminal space of some lesions contained yellow appearing material, indicative of a mixture of mucus and surfactant secretions. Interestingly, we also detected cystic lesions that were lined by a metaplastic epithelium characterized by an atypical transition from a SFTPC^+^ epithelium to a MUC5B^+^ epithelium in all analyzed IPF samples (**Figure 8G, Figure S9**). These transition zones of ATII-like cells residing in close proximity to goblet-like cells are indicative of a loss of regional epithelial specification and could represent areas of progressing alveolar bronchiolization.

## DISCUSSION

In this study, we establish a novel iPSC-derived ALI model of alveolar epithelial differentiation to investigate epithelial dysfunction related to IPF. Utilizing this system, we demonstrate how an IPF-relevant cytokine environment can skew iPSC-derived ATII-like differentiation towards proximal lineages and induce disease-related changes, including secretion of IPF biomarkers. This report thereby describes, for the first time, a human model system that recapitulates key aspects of IPF-related bronchiolization *in vitro*.

While ALI culture has been applied to human pluripotent stem cell (hPSC)-derived airway epithelial cells to enhance maturation, this method has not yet been tested in hPSC-derived models of the alveolar epithelium (49, 50). We show for the first time, the maturation of hPSC-derived lung epithelial progenitor cells towards ATII-like cells in an ALI culture format, providing a novel platform for both basic research and drug discovery purposes. Our data show that iPSC-derived lung epithelial progenitor cells differentiated at ALI express higher levels of *SFTPC, SFTPB* and *ABCA3* compared to submerged controls. Importantly, SFTPC^+^ cells are located on the apical surface of the airlifted cultures, suggestive of a positive influence of air exposure on ATII-like differentiation. Interestingly, ABCA3 positive cells, which are not detectable in human fetal lungs prior to 22-23 weeks of gestation, are present within our system (51). The cultures also express *TM4SF1* which has been described as a marker for alveolar epithelial progenitor cells within the adult human lung (16). In addition, lamellar bodies, ATII specific surfactant producing/storing organelles, are present in the cultures (28, 52-54). Taken together, these results confirm the feasibility of ATII-like cell derivation from iPSC-derived lung progenitor cells in ALI culture.

The disease specific milieu within the lung has been suggested to play a central role in the pathogenesis of IPF (55, 56). While there is substantial evidence for the relevance of various cytokines and growth factors in IPF, including TGF-β, TGF-α, IL-13, TNF-α and IL-1β, the overexpression of which induces pulmonary fibrosis in animal models, no single factor is known to simultaneously activate all IPF-related pathways (57, 58). Despite this fact, many currently available *in vitro* models of pulmonary fibrosis rely on a single stimulus (59). To mimic a pro-fibrotic milieu *in vitro*, we designed a novel cocktail (IPF-RC) containing nine IPF-relevant cytokines and assessed its effect on differentiating iPSC-derived distal lung epithelial progenitor cells (38-46). As we primarily aimed to investigate the effect of a fibrosis-relevant environment on progenitors of the alveolar epithelium, IPF-RC stimulation was initiated by day 35 of the protocol, following induction of *SFTPC* expression, which constitutes a specific feature of alveolar epithelial progenitors in fetal human lungs (60-62).

Analysis of the culture medium in IPF-RC treated cells showed increased MMP-7 and MMP-10, both of which are elevated in IPF patient BAL and serum, and RNA-seq analysis revealed an enrichment of transcripts involved in extracellular matrix organization, indicating an induction of IPF-relevant processes (63-65). Furthermore, a comparison of transcriptional changes in IPF-RC treated iPSC-derived cultures and human IPF patient lungs revealed a significant overlap of up- and down-regulated transcripts (48). On the other hand, there is also a substantial amount of non-overlapping deregulated genes, likely both due to the heterogeneity of our culture system and the origin of the publicly available IPF patient dataset. The analysis of whole human lung homogenates contains multiple cell types (e.g. mesenchymal, endothelial and immune cells) that are not represented in our *in vitro* model system. This is supported by the presence of known epithelial-specific transcripts among the commonly deregulated genes, many of which have been associated with lung fibrosis. In accordance with previous reports, *KLF4, VEGFA, IL32* and *SLC19A3* are among the commonly downregulated transcripts (7, 66-68). *BPIFB1* which was described to localize to the bronchiolized epithelium in the honeycomb cysts in usual interstitial pneumonia and *MMP10*, are among the commonly up-regulated transcripts (69, 70). Taken together, our iPSC-derived *in vitro* model mimics certain IPF-related changes in epithelial transcription and secretory phenotype.

In addition, RNA-seq data reveal many known airway associated transcripts with known roles in the developing and adult proximal lung, were among the top upregulated transcripts in IPF-RC treated iPSC-derived cultures, e.g. *SOX2, PAX9, TP63, KRT5, FOXJ1, SCGB1A1* and *MUC5B*, which have recently been confirmed as markers for the altered IPF lung epithelium via single-cell sequencing (5, 7, 9, 70-72). Upregulation of airway marker expression following IPF-RC stimulation is accompanied by a loss of ATII specific *SFTPC* expression and a shift in the expression of *SOX2* and *SOX9*, two transcription factors known as central players in proximo-distal epithelial patterning during human and mouse lung development (28, 61). Mouse studies have revealed an essential role of Sox9 during branching morphogenesis, correct distal lung epithelial differentiation during development, as well as recovery of lung function after acute lung injury in adult mice (73, 74). Expression of SOX2, which is associated with proximal airway rather than alveolar identity in healthy lungs, has been shown in the bronchiolized and enlarged distal airspaces in IPF (5). Moreover, two murine studies have independently shown that selective overexpression of *Sox2* in adult ATII cells resulted in an induction of conducting airway-related transcript expression in the alveoli (75, 76). Immunohistological analysis of iPSC-derived cultures differentiated in the presence of IPF-RC revealed a significant reduction in SFTPC^+^ cell area and the emergence MUC5B^+^ secretory-like cells, an important role of which has been suggested in IPF.

Not only has it been shown that MUC5B^+^ cells abundantly reside within the bronchiolized IPF epithelium, but a *MUC5B* promoter polymorphism, known to enhance MUC5B expression in the bronchiolo-alveolar epithelia, constitutes the strongest currently known genetic risk factor for IPF (5, 6, 77, 78). We further demonstrated in this study, that highly aberrant direct transition zones between SFTPC^+^ and MUC5B^+^ epithelium exist within distal IPF patient lungs. Based on our findings in a human iPSC-derived alveolar epithelial model and IPF patient lungs, we hypothesize that alveolar epithelial progenitor cells could contribute to epithelial bronchiolization of the distal compartments of the IPF lung via aberrant trans-differentiation towards airway-like lineages.

In our model system, removal of IL-13, but not of TGF-β1 or TNF-α, from IPF-RC impairs its ability to promote differentiation towards airway-like cell fates at the cost of ATII-like cell differentiation, indicating IL-13 as one of the main drivers of the proximalization effect we observed following IPF-RC stimulation. IL-13 has been described as a main mediator in respiratory diseases, particularly in asthma, where it is linked to mucus secretion and fibrogenic processes (79). Moreover, IL-13 overexpression leads to pulmonary fibrosis in mice and it has been hypothesized that IL-13 antagonism could constitute a potential treatment strategy for IPF (80). While IL-13 alone is sufficient to repress *SFTPC* expression, in line with the previous findings in primary human ATII cells (81), it fails to fully recapitulate the influence of the IPF-RC on airway-specific and IPF-relevant transcripts. Our results are in accordance with the current consensus that neither IL-13, nor any other single factor is sufficient to simultaneously activate all IPF-related pathways *in vitro* and highlight the importance of more physiological human *in vitro* model systems (58). This is underlined by the recent failure of the phase 2 clinical study of tralokinumab, a human anti-IL-13 monoclonal antibody, in subjects with IPF (82).

It has been proposed that migrating airway stem cells, rather than resident alveolar stem cells, are the stem cell source of the bronchiolized epithelium. As our model system contains rare individual KRT5^+^ basal-like cells, their role in the IPF-RC mediated effects cannot be excluded. However, as IPF-RC treatment of small airway basal cells induced a strong upregulation of *MUC5AC*, but not *MUC5B*, contrary to the abundant emergence of aberrant MUC5B^+^ cells in the bronchiolized distal airspaces in IPF patients, as well as suppressed the expression of transcripts upregulated in the IPF lung such as SCGB1A1 and FOXJ1, the airway-like phenotype observed in iPSC-derived IPF-RC cultures is unlikely the sole result of basal-like cell expansion and differentiation (5, 6, 70, 71).

The usual interstitial pneumonia pattern in IPF is characterized by high temporal and spatial heterogeneity (83). We hypothesize that our *in vitro* system utilizing an IPF-relevant cytokine cocktail, could potentially model the progression of epithelial dysfunction from severely affected towards healthy neighboring regions of the lung via a distribution of the pro-fibrotic milieu produced in diseased fibrotic areas within the lung and exposure of epithelial progenitor cells in previously unaffected regions to this milieu. However, as the definite composition of the IPF BAL/lung milieu is not known, the combination of nine cytokines cannot fully mimic the environment epithelial cells are exposed to in a diseased lung. Moreover, the model utilized in this study does not encapsulate the complex cellular interplay that leads to the vicious cycle of wound repair and scar formation that occurs in IPF (84), nor the role of genetic risk factors in IPF pathogenesis (18).

Overall, our findings highlight that the fibrosis-related milieu present in the IPF lung has the potential to skew alveolar epithelial differentiation towards airway cell phenotypes. Our data suggest that aberrant trans-differentiation of epithelial stem cells in the fibrotic lung could disrupt the regular proximo-distal patterning of the lung epithelium and thereby contribute to the emergence of aberrant epithelial cell types, as well as the apparent bronchiolization in the distal IPF lung. This raises the need for further investigations to determine the exact pathophysiological mechanisms underlying alveolar epithelial cell dysfunction in IPF, which may open routes to novel therapeutic concepts not yet covered by current standard of care therapies.

## Supporting information

Supplementary RNA-seq data

