## Supplementary Information for "Recapitulating idiopathic pulmonary fibrosis related alveolar epithelial dysfunction in an iPSC-derived air-liquid interface model"

**This PDF file includes:**

SI Materials and Methods

Figures S1 to S9

Tables S1 to S3

References for SI reference citations

### SI MATERIALS AND METHODS

#### *Composition of an IPF-relevant cytokine cocktail (IPF-RC)*

An IPF-relevant cytokine cocktail (IPF-RC) was designed based on literature studies that compared both IPF and healthy control bronchoalveolar lavage or sputum samples (

**Table** S1). Nine cytokines were selected, based on being significantly upregulated in IPF samples compared to healthy controls. To approximately account for dilution in saline during sample collection, cytokine concentrations one order of magnitude higher than measured in IPF patient BAL were estimated as suitable for *in vitro* experiments. Long-term stimulation of cells with IPF-RC was performed to simulate the cytokine milieu present in an IPF lung. Final human recombinant cytokine concentrations (all acquired from R&D Systems, Minneapolis, MN, US) in the cell culture medium during IPF-RC treatment were 300 pg/ mL TGF-β1, 10 pg/ mL IL-1β, 100 pg/ mL TNF-α, 1500 pg/ mL IL-8, 700 pg/ mL MCP1, 40 pg/ mL IL-33, 100 pg/ mL TSLP, 2500 pg/ mL IL-13 and 160 pg/ mL IL-4, respectively. IPF-RC was prepared as a 1000x stock in 0.1% BSA in PBS.

#### *IPF patient samples*

Formalin-fixed and paraffin embedded lung samples from human IPF patients were purchased from Folio Biosciences (Powell, OH, US) under the regulatory conditions of the Boehringer Ingelheim corporate policy regarding the acquisition and use of human biospecimen. Samples were reviewed internally by a trained pathologist (on H&E basis) and the initial diagnosis of IPF was confirmed.

#### *Human induced pluripotent stem cell culture*

The human iPSC lines utilized in this study were obtained from the StemBANCC consortium and were reprogrammed from human fibroblasts via non-integrating sendai virus. iPSC lines

SFC065-03-03 (EBiSC: STBCi057-A, Biosamples ID: SAMEA104493762) and SFC084-03-01 (EBiSC: STBCi033-A, Biosamples ID: SAMEA104493681) were cultured in mTeSR 1 (Stemcell Technologies, Vancouver, Canada) on hESC-qualified matrigel (Corning, New York, US) coated cell culture plates at 37 °C/ 5% $CO_2$ in a humidified normoxic incubator and passaged using 0.5 mM EDTA pH 8.0.

***Differentiation of human iPSCs into lung epithelial progenitor cells***

Directed differentiation of iPSCs towards NKX2.1$^+$ lung progenitor cells was performed in serum-free differentiation (SFD$^+$) medium, as previously described (1, 2). Briefly, definitive endoderm (DE) formation of embryoid bodies was induced in the presence of Activin A, followed by dissociation and replating and anterior foregut endoderm (AFE) induction by BMP, TGF-β and Wnt inhibition. Ventralization was achieved by applying Wnt, BMP, FGF and RA signaling. After 14 days of differentiation, VAFE cells were detached as clumps and either frozen down for later usage or replated and expanded for 10 additional days on hESC-qualified matrigel (Corning, New York, US) coated cell culture plates.

***Branching organoid formation of iPSC derived lung epithelial progenitor cells***

Branching of iPSC derived lung epithelial progenitor cells was induced in 3D culture, similar to previous reports (3). Cells were cultured in SFD$^+$ medium supplemented with 3 µM CHIR99021 (Axon Medchem, Groningen, Netherlands), 10 ng/ mL rhKGF (R&D Systems, Minneapolis, MN, US) and 10 ng/ mL rhFGF10 (R&D Systems, Minneapolis, MN, US). On day 24 of differentiation, lung progenitor cells were briefly digested with warm 0.05% Trypsin/ 0.53 mM EDTA and detached as clumps, which were collected at the bottom of a canonical tube via sedimentation and plated onto low attachment dishes. The following day, the resulting organoids were embedded into cold growth factor reduced matrigel (Corning, New York, US) on 48-well tissue culture plates. Following solidification of the matrigel, the organoids were covered with medium and cultured for up to 35 days. The medium was changed every two days.

***iPSC derived lung epithelial progenitor maturation into ATII-like cells in 2D air-liquid interface culture***

On day 24 of differentiation, lung progenitor cells were briefly digested with warm 0.05% Trypsin/ 0.53 mM EDTA and detached as clumps and collected via sedimentation. The cells were plated onto hESC-qualified matrigel (Corning, New York, US) coated transwell permeable support inserts (PET membrane, 12 well format, pore size 0.4 µm, Corning, New York, US) and cultured under submerged conditions until day 32 to allow the cells to attach and to spread out over the whole insert surface. Subsequently, the apical medium was removed and the cells were cultured at air-liquid interface until day 49 of differentiation. Similar to a previous study, temporal withdrawal of Wnt signaling was employed to promote distal lung epithelial progenitor differentiation towards ATII-like cells (4). From day 24 to day 35 of differentiation, lung progenitors were cultured in $SFD^+$ medium supplemented with 3 µM CHIR99021 (Axon Medchem, Groningen, Netherlands), 10 ng/ mL rhKGF (R&D Systems, Minneapolis, MN, US), 10 ng/ mL rhFGF10 (R&D Systems, Minneapolis, MN, US), 25 ng/ mL dexamethasone (Sigma-Aldrich, St. Louis, MO, US), 0.1 mM 8-Br-cAMP (Sigma-Aldrich, St. Louis, MO, US) and 0.1 mM 3-Isobutyl-1-methylxanthine (Sigma-Aldrich, St. Louis, MO, US). Subsequently, the cells were treated with the same medium, but without CHIR99021, from day 35 to 42, before 3 µM CHIR99021 was added back for one more week until day 49 of differentiation. The culture medium was exchanged every other day during the whole maturation phase. Where indicated in the text, additional treatments were applied during differentiation from day 35 to day 49.

***ATI-like differentiation of iPSC derived ATII-like cells***

ATI-like differentiation of iPSC derived ATII cells was induced by replating cells on plastic in 2D submerged culture, as previously described (4). On day 49 of differentiation ATII-like cells were dissociated using Gibco StemPro Accutase cell dissociation reagent (Life Technologies.

Carlsbad, CA, US) and replated onto 6-well tissue culture plates in Gibco DMEM Glutamax (Life Technologies. Carlsbad, CA, US) supplemented with 10% FBS (Life Technologies. Carlsbad, CA, US). Media was changed every other day and cells were harvested for analysis after five days. Controls were not replated and remained at air-liquid interface on transwell permeable support inserts until they were harvested on day 54 of differentiation.

***Primary human lung fibroblast culture***

Primary human lung fibroblasts (CC-2512, Lot. 0000608197, Lonza, Basel, Switzerland) were grown in fibroblast basal medium (FBM) (Lonza, Basel, Switzerland) supplemented with FGM-2 SingleQuot Kit Supplements & Growth Factors (Lonza, Basel, Switzerland) at 37 °C and 5% $CO_2$. Cells were passaged maximum 10 times before use.

***Primary human small airway epithelial cell culture***

Primary human Small Airway Epithelial Cells (SAECs) (CC-2547, Lot 501937, Lonza, Basel, Switzerland) were thawed into a T-175 tissue culture flask and expanded for four days in PneumaCult-Ex Plus Medium (Stemcell Technologies, Vancouver, Canada). A single cell suspension was then created using an Animal Component-Free Cell Dissociation Kit (Stemcell Technologies, Vancouver, Canada) and 90000 cells/ $cm^2$ were seeded onto rat tail collagen type I (Corning, New York, US) coated transwell permeable support inserts (PET membrane, 24 well format, pore size 0.4 µm, Corning, New York, US). Four days post seeding, the apical medium was removed and the cells were differentiated at air-liquid interface (ALI) in PneumaCult-ALI 2 medium (Stemcell Technologies, Vancouver, Canada) for 28 days. The culture medium was exchanged every other day. Where indicated in the text, IPF-RC treatment was applied to SAEC cultures from 1 to 28 days post-ALI.

***Quantitative Real-Time PCR***

Cells were lysed in RLT Plus buffer (Qiagen) and RNA isolation was performed using the RNeasy Plus Mini Kit (Qiagen, Venlo, Netherlands) according to the manufacturer's instructions. In some experiments, human lung total RNA (AM7968, Lot.1850512, Invitrogen, Life Technologies. Carlsbad, CA, US) was used as a positive control. Reverse transcription was performed using the Applied Biosystems High-Capacity cDNA Reverse Transcription Kit (Life Technologies. Carlsbad, CA, US) according to the manufacturer's instructions. 2µl (5 ng) of cDNA were added to a final reaction volume of 10 µl containing the QuantiFast Probe PCR +ROX Vial Kit (Qiagen, Venlo, Netherlands) master mix and the respective Applied Biosystems TaqMan Gene Expression Assay FAM (Life Technologies. Carlsbad, CA, US). qPCR was performed in 384-format using an Applied Biosystems ABI ViiA 7 real-time PCR System (Life Technologies. Carlsbad, CA, US). Gene expression was normalized to GAPDH control and fold change in gene expression relative to iPSCs (day 0 of differentiation) was calculated using the $2^{(-\Delta\Delta CT)}$ method. A table of all TaqMan Gene Expression Assays used in this study is provided in the Supplementary Information (SI Appendix, **Table S3**)

***RNA extraction, Illumina library preparation and RNA sequencing***

Total RNA of iPSC-derived day 0, day 24 and day 49 cultures (three biological replicates per timepoint) was extracted using the Ambion Magmax-96 total RNA isolation kit (Life Technologies. Carlsbad, CA, US) according to the manufacturer's instructions. Nucleic acids were captured onto magnetic beads, treated with DNase and total RNA was eluted in 50 µl elution buffer. RNA quality and concentration were measured using an RNA Pico chip on a Bioanalyzer (Agilent, Santa Clara, CA, US).

A sequencing library was prepared using the TrueSeq RNA Sample Prep Kit v2-Set B (Illumina, San Diego, CA, US) with 200 ng of total RNA input per sample, resulting in an average fragment size of 275 bp including adapters. Before sequencing, eight individual libraries were normalized and pooled together using the adapter indices supplied by the manufacturer. Pooled libraries were then clustered on an Illumina cBot Instrument using the TruSeq SR Cluster Kit v3—cBot—HS (Illumina, San Diego, CA, US). Sequencing was performed as 50 bp, single reads and 7 bases index read on an Illumina HiSeq2000 instrument using the TruSeq SBS Kit HS- v3 (50-cycle) (Illumina, San Diego, CA, US).

#### *Bioinformatic analysis of RNA sequencing data*

Step-by-step RNA sequencing pipeline

A step-by-step bioinformatics pipeline was utilized to process and technically validate the RNA sequencing data. The pipeline was described in detail in a previous study (5).

Detection of differentially expressed genes

Comparative analysis was done using the limma R-package (6). Benjamini-Hochberg correction was used to adjust for multiple testing.

Heatmaps

A heatmap of the top 500 genes deregulated in iPSC-derived day 49 cultures versus day 24 cultures was created utilizing the heatmap3 package of R statistical software and the following cut-off values: q-val < 0.001, |logFC| > 2, maxRPKM > 10.

Principle component analysis (PCA)

PCA was performed as previously described, using ClustVis (7). The top 5000 transcripts according to the coefficient of variation were selected for analysis.

Gene set enrichment analysis (GSEA)

Normalized expression data for day 49 +IPF-RC and day 49 samples were analyzed using the GSEA method (8). Genes were ranked by magnitude of correlation with a class distinction, and the GSEA algorithm determined whether members of a gene set tended to occur towards the top or bottom of the list, and calculated an enrichment score. In order to remove extremely low or not at all expressed transcripts from the data set, NGS data was filtered according to an RPKM value > 2 in at least one of the samples.  Gene sets that were significantly associated with the day 49 +IPF-RC class were identified from the C2.CP.REACTOME (Curated) collection of gene sets in the Molecular Signatures Database v6.2 (MSigDB), which comprises

674 gene sets (9). The null distributions used to calculate the statistical significance of the enrichment scores were generated by permutation of gene sets for the day 49 +IPF-RC vs. day 49 contrast (3 replicates/class). Adjustment for multiple hypothesis testing was performed by determination of FDR. Enrichment scores with an FDR q-value <0.05 were considered to be significant.

<u>Calculation of intersections with IPF lung RNA sequencing dataset</u>

Expression data for day 49 +IPF-RC and day 49 samples were compared to publicly available data for human IPF and healthy lung tissue (10). Transcripts used for calculating the intersections were selected according to being expressed (at least one sample showing an RPKM >2) in both data sets. This resulted in a universe of 12365 transcripts. De-regulated transcripts were defined by exhibiting a q-value <0.05 and an absolute log ratio of >0.5. Resulting gene sets were split into up- and down-regulated and the intersection of the corresponding gene sets from both data sets was calculated. Significance of the intersection was tested via a hypergeometric test.

***Measurement of total cell number and Caspase-3/ 7 activity***

The total cell number per insert of iPSC-derived day 49 cultures was assessed by dissociating the cultures with Gibco StemPro Accutase cell dissociation reagent (Life Technologies. Carlsbad, CA, US) and counting of cells with a TC20 Automated Cell Counter (Bio-Rad, Hercules, CA, US). Caspase-3/ 7 activity of day 49 cultures was measured using a Caspase-Glo® 3/7 Assay Kit (Promega, Madison, WI, US) according to the manufacturer’s instructions.

***Measurement of MMP protein concentrations by ELISA***

MMP-7 and MMP-10 protein concentrations in cell culture supernatants were determined using a Human Total MMP-7 DuoSet ELISA (R&D Systems, Minneapolis, MN, US) and a Human Total MMP-10 DuoSet ELISA (R&D Systems, Minneapolis, MN, US) according to the manufacturer’s instructions.

***Flow cytometry***

FACS was performed on day 24 of differentiation. Lung progenitors were dissociated using Gibco StemPro Accutase cell dissociation reagent (Life Technologies. Carlsbad, CA, US) and taken up into an excess of ice cold 1% BSA in PBS. All subsequent washing steps were performed by applying ice cold wash buffer (0.1% BSA in PBS) and pelleting at 300g for 5min. 1*10^6 cells were transferred into a FACS tube and washed, before prewarmed Cytofix fixation buffer (BD Biosciences, Franklin Lakes, NJ, US) was added and the sample was incubated for 10 min at 37 °C. Cells were then washed and permeabilized in 80% MeOH for 5 min, followed by 0.1% Tween-20 (Sigma-Aldrich, St. Louis, MO, US) in wash buffer for 10 min. Blocking was performed in 10% goat serum (Sigma-Aldrich, St. Louis, MO, US) in PBS for 10 min. Cells were washed and incubated in the respective antibody solution for 30 min in the dark. Unstained controls were incubated in wash buffer, isotype controls were incubated in 1 µg/ mL Alexa Fluor 488 conjugated rabbit IgG Isotype Control (Abcam, Cambridge, UK) and samples for NKX2.1 staining were incubated in 1 µg/ mL Alexa Fluor conjugated anti-TTF1 antibody (Abcam, Cambridge UK). The cells were then washed, resupended in Cellfix (BD Biosciences) and transferred into fresh FACS tubes through a 35 µm strainer cap. Samples were measured on a LSR II flow cytometer (BD Biosciences, Franklin Lakes, NJ, US) and analysed using the FACSDiva (BD Biosciences, Franklin Lakes, NJ, US) and the FloJo (FlowJo LLC) software.

***Immunohistochemistry***

A table of all primary and secondary antibodies used in this study is provided in the Supplementary Information (SI Appendix, **Table S2**).

Haematoxylin and eosin and Alcian Blue/ PAS stainings

Transwell cultures were fixed for 30 minutes with 4 % Paraformaldehyde (BosterBio, Pleasanton, CA, US) at room temperature and the whole membrane was cut out of the insert for further processing. Fixed inserts were dehydrated and embedded into paraffin following standard procedures. Transwell insert cross-sections or sections of paraffin-embedded IPF lung samples of 3 µm thickness were prepared and rehydrated using a descending series of ethanol. Haematoxylin and eosin (H&E) and Alcian Blue/ PAS stainings were performed according to standard protocols. Images were acquired with an AxioCam MR3 (Zeiss, Oberkochen, Germany) using the 20x objective of an Axio Imager Z1 (Zeiss, Oberkochen, Germany).

Immunofluorescence of iPSC-derived and primary cells

For immunofluorescence of lung epithelial progenitor cells, VAFE cell clumps were replated and expanded on hESC-qualified matrigel (Corning, New York, US) coated glass bottom plates (12-well, No. 1.5 coverslip, 14 mm glass diameter, MatTek, Ashland, MA, US). On day 24 of differentiation, cells were fixed for 15 min with 4 % Paraformaldehyde (BosterBio, Pleasanton, CA, US) at room temperature. The fixed samples were washed with PBS and blocked for one hour at room temperature in 5% BSA with 0.3% TritonX-100 (Sigma-Aldrich, St. Louis, MO, US) in PBS. The cells were then incubated with the respective primary antibodies diluted in antibody dilution solution (DCS, Hamburg, Germany) over night at 4 °C. On the following day, the samples were washed with PBS and incubated with the respective fluorophore conjugated secondary antibodies, diluted 1:500 in antibody dilution solution, for one hour at room temperature. To visualize the nuclei, the cells were then stained with 2 µg/ mL Hoechst 33342 (Life Technologies. Carlsbad, CA, US) in PBS for 5 min and washed with PBS. Subsequently, the finished samples were then covered with 1 mL PBS and imaged.

For immunofluorescence of transwell inserts, iPSC derived ATII-like cells or primary SAEC cultures were fixed for 30 minutes with 4 % Paraformaldehyde (BosterBio, Pleasanton, CA, US) at room temperature. The inserts were then washed with PBS and the whole membrane was cut out of the insert for further processing.

Whole insert stainings were performed after blocking the fixed cells on the membrane for one hour at room temperature in 5% BSA with 0.3% TritonX-100 (Sigma-Aldrich, St. Louis, MO,

US) in PBS, by incubating with primary antibodies diluted in antibody dilution solution (DCS, Hamburg, Germany) over night at 4 °C. After washing, the samples were incubated with the respective fluorophore conjugated secondary antibodies, diluted 1:500 in antibody dilution solution for one hour at room temperature. The membranes were then washed in PBS, mounted with Invitrogen ProLong Diamond Antifade Mountant with DAPI (Life Technologies. Carlsbad, CA, US), coverslipped and imaged.

For staining of insert cross-sections and definitive endoderm bodies, fixed specimen were dehydrated using a Tissue Tek VIP processor and embedded into paraffin blocks. Sections of 3 µm thickness were prepared, deparaffinized with xylene and rehydrated using a descending series of ethanol. Antigen retrieval was performed for 30 min at 95 °C in H.I.E.R. sodium citrate solution (BioLegend, San Diego, CA, US). The samples were blocked for one hour at room temperature with 5% normal goat or normal donkey serum (Sigma-Aldrich, St. Louis, MO, US) in PBS and incubated over night at 4 °C with the respective primary antibodies diluted in antibody dilution solution (DCS, Hamburg, Germany). The samples were washed in PBS with 0.5% Tween-20 (Sigma-Aldrich, St. Louis, MO, US) and incubated with the respective fluorophore conjugated secondary antibodies, diluted 1:500 in antibody dilution solution, for one hour at room temperature. Following washing of the slides with PBS with 0.5% Tween-20 (Sigma-Aldrich, St. Louis, MO, US), the sections were mounted with Invitrogen ProLong Diamond Antifade Mountant with DAPI (Life Technologies. Carlsbad, CA, US) and coverslipped.

All specimens were imaged with a LSM 710 confocal microscope system (Zeiss, Oberkochen, Germany) with an AXIO Observer Z1 using a Plan-Apochromat 20x objective or a Plan-Apochromat 40x water-immersion objective.

Immunohistochemistry of IPF lung sections

Immunohistochemistry in IPF patient samples was carried out on the automated Leica IHC Bond-RX platform (Leica Biosystems, Nussloch, Germany) using the Opal method (Perkin Elmer, Waltham, MA, US). Three µm thick sections of FFPE IPF lung tissue on super frost plus slides were deparaffinised and rehydrated for multiplex immunohistochemistry staining. Antigen retrieval was performed for all primary antibodies by heating the sections in Bond ER solution 1 buffer (Leica Biosystems, Nussloch, Germany) at 95°C; pH 6.0 for 20 min. Primary antibodies were diluted with Leica Primary Antibody Diluent (Leica Biosystems, Nussloch, Germany) and sections were incubated with them for 30 min at room temperature followed by incubation (10 min at RT) with Opal Polymer Anti-Rabbit HRP Kit (ARR1001KT; Perkin Elmer Waltham, MA, US). Immunofluorescent signal was visualized using the OPAL (Perkin Elmer, Waltham, MA, US) TSA dye 570 and TSA dye 650 and counterstained with Spectral DAPI (Akoya, Menlo Park, CA). Microscopy of IPF lung samples was conducted with an AxioImager M2 microscope (Zeiss, Oberkochen, Germany) and images were created using an AxioScan scanner and ZEN slidescan software (Zeiss, Oberkochen, Germany).

***Semi-quantitative measurement of SFTPC$^{+}$ area in iPSC-derived cultures***

Immunofluorescence of fixed whole transwell inserts was performed as described above. All samples of an experiment were stained and imaged on the same day. Nine randomly selected areas per insert (425.1 µm x 425.1 µm per area) were captured using a 20x Plan-Apochromat objective on a LSM 710 confocal microscope system (Zeiss, Oberkochen, Germany) with the following settings: Alexa Fluor 488 channel, 5% laser power, master gain 600, acquisition speed 9, resolution 1024 px x 1024 px. To account for unevenness in specimen surface and thickness, Z-stack imaging of 20 vertical stacks per area was applied and maximum intensity projection was performed using the ZEN 2012 Black Edition software (Zeiss, Oberkochen, Germany). Image analysis was then performed using the Fiji for ImageJ software (11). Each image was first converted to 8-bit format and then converted to binary at a threshold of 20-255. In the following, particles within areas >80 px were analysed and summarized per image. For each analyzed transwell insert, the mean positive area per image was calculated from the nine selected areas.

***Transmission electron microscopy***

For electron microscopy of iPSC derived ATII-like cells, transwell inserts were pre-fixed for 1 h with 4 % Paraformaldehyde. The PET membrane with pre-fixed cells was then washed with a 0,1% cacodylate buffer solution for 30 minutes. Afterwards, the samples were transferred into a EM-TP Tissue processor (Leica Microsystems, Wetzlar, Germany) for automated post-fixation, staining, dehydration and embedding. In detail, the samples underwent the following steps: 20 min in 0,1% cacodylate buffer, 3 h in 1% Daltons osmiumtetroxide aq., 3 times 15 min in 0,1% cacodylate buffer solution, 15 min in 30% isopropanol, 30 min each in 30%, 50%, 70%, 90% and 100% isopropanol, and three times 1 h in 100% isopropanol. Following dehydration, sample infiltration with Epoxy resin was achieved as follows: 30 min in 50% isopropanol/ 50% EPON, 30 min in 33% isopropanol/ 66% EPON, 30 min in 20% isopropanol/ 80% EPON and 60 min in 100% EPON. The samples were then incubated twice for 6 h in 100% EPON and hardened at 60°C for 24 hours. Ultra-thin sections (50 nm) were prepared on an Ultracut UCT ultra-microtome (Leica Microsystems, Wetzlar, Germany) and imaged on a TEM 912AB (Zeiss, Oberkochen,Germany).

***LysoTracker Green DND-26 live cell staining***

iPSC derived ATII-like cells were dissociated using Gibco StemPro Accutase cell dissociation reagent (Life Technologies. Carlsbad, CA, US) and replated onto hESC-qualified matrigel (Corning, New York, US) coated Nunc 8-well Lab-Tek Chambered Coverglasses (Thermo Fisher Scientific, Waltham, MA, US) in $SFD^+$ medium, supplemented with 3 µM CHIR99021 (Axon Medchem, Groningen, Netherlands), 10 ng/ mL rhKGF (R&D Systems, Minneapolis,

MN, US), 10 ng/ mL rhFGF10 (R&D Systems, Minneapolis, MN, US), 25 ng/ mL dexamethasone (Sigma-Aldrich, St. Louis, MO, US), 0.1 mM 8-Br-cAMP (Sigma-Aldrich, St. Louis, MO, US), 0.1 mM 3-Isobutyl-1-methylxanthine (Sigma-Aldrich, St. Louis, MO, US) and 10 µM Y-27632 (Abcam, Cambridge, UK). The cells were allowed to attach overnight, before 100 nM LysoTracker Green DND-26 (Invitrogen, Life Technologies. Carlsbad, CA, US) was added into the medium and the cells were incubated for 20 min at 37 °C protected from light. The medium was then removed and the samples were washed once with prewarmed Gibco FluoroBrite DMEM (Life Technologies. Carlsbad, CA, US), followed by a 5 min incubation in 5 ng/ mL Hoechst 33342 (Life Technologies. Carlsbad, CA, US) in FluoroBrite. Subsequent to three additional washes with FluoroBrite DMEM, fresh FluoroBrite DMEM was added to the cells and the samples were immediately imaged using a Plan-Apochromat 63x oil-immersion objective on an LSM 710 confocal microscope system (Zeiss, Oberkochen, Germany).

***Visual collagen I quantification in primary human lung fibroblast cultures***

Primary human lung fibroblasts were plated in a poly-D-lysine coated 384 CellCarrier microtiter plate from PerkinElmer in fibroblast basal medium (FBM) with FGM-2TM Single Quots (Lonza, Basel, Switzerland) at a density of 1000 cells per well. After 24 h, the medium was replaced by the same medium containing no serum (starvation medium). 48 h after cell seeding, the starvation medium was replaced with starvation medium containing a mixture of Ficoll 70 and 400 (GE Healthcare, Chicago, IL, US; 37.5 mg/ mL and 25 mg/ mL, respectively), 200 µM vitamin C and IPF-RC (1:1000 dilution). After 72 h, the cell culture medium was removed and cells were fixed with 100% ice-cold methanol for 30 minutes. Next, cells were washed with PBS, permeabilized for 20 minutes using 1% Triton-X-100 (Sigma-Aldrich, St. Louis, MO, US), washed and blocked for 30 minutes with 3% BSA in PBS. After an additional wash step, cell nuclei were stained with 1 µM Hoechst 33342 (Life Technologies. Carlsbad, CA, US) and collagen I was stained using a monoclonal antibody (SAB4200678, Sigma-

Aldrich, St. Louis, MO, US). For primary antibody detection, cells were washed and incubated for 30 minutes at 37 °C with Goat anti mouse IgG1 Alexa Fluor 568 secondary antibody. After secondary antibody removal, cells were stained with Invitrogen HCS Cell Mask Green stain (1:50000, Life Technologies. Carlsbad, CA, US). Following a final wash step, images were acquired in an InCell 2200 Analyzer (GE Heathcare, Chicago, IL, US), using 2D-Deconvolution for nuclei (Hoechst channel), cells (FITC channel), and collagen I (TexasRed channel), and images were transferred to the Columbus Image Storage and Analysis system (Perkin Elmer, Waltham, MA, US).

Image analysis was performed as previously described (12, 13). Briefly, using the building blocks of Perkin Elmer´s Columbus Image Analysis system, first nuclei acquired with the Hoechst channel were detected using the building block (BB) "nuclei". Second, cells were defined with the BB "find cytoplasm" from the FITC channel. Collagen I area was defined by two individual BBs "find simple image region" based on images acquired in the TexasRed channel. Collagen I readouts were normalized to the number of cells per image field. Total number of cells, and total collagen area/total number of cells were used as parameters to quantify effects of IPF-RC.

***Cilia beat measurements in primary small airway cultures***

Cilia beat frequency was calculated from image stacks of 2D+time. Six evenly distributed regions of a transwell insert were imaged using the 32x objective of an Axiovert 25 microscope (Zeiss, Oberkochen, Germany) and an acA 1300-200µm black and white USB-3.0 high speed camera (Basler, Ahrensburg, Germany). Ciliary movement was recorded at 100 frames per second for a total of 6 seconds per region. Applications for image capture and analysis were developed using the Halcon 13.0.2 machine vision software toolbox (MVTec Software, Munich, Germany). Visualization of image stack was performed with Analyze (AnalyzeDirect, Overland Park, KS, US). A grey value time course was calculated for each pixel over 512 frames per region. The area covered by motile cilia was determined by quantifying the percentage of pixels with a measurable beating frequency within a region. The mean ciliary beating frequency of a region was determined by calculating the average frequency of the change in grey values.

***Statistical analysis***

All qRT-PCR, ELISA, FACS and image analysis data are presented as mean with error bars representing the SEM of at least three independent experiments. Experiments comparing two conditions only were analyzed using two-tailed unpaired Student's t-test, when variances were homogeneously distributed, or Mann-Whitney U-test, when data were not normally distributed. Experiments containing three or more conditions were assessed by two-way ANOVA, followed by the Tukey test. Data analysis was performed using GraphPad Prism 8.0 (GraphPad Software). Significance values are marked as * $P < 0.05$, ** $P < 0.01$, *** $P < 0.001$ and **** $P < 0.0001$.

### SI FIGURES

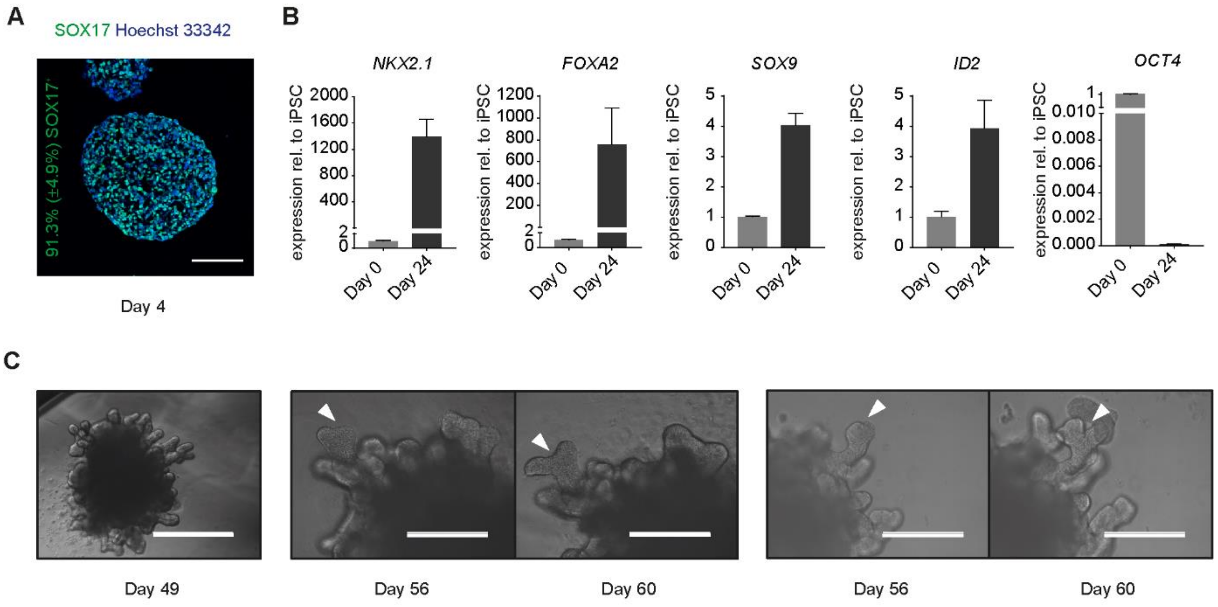


**Figure S1. Directed differentiation of the human iPSC line SFC065-03-03 towards VAFE cells displaying expression of distal lung epithelial progenitor cell markers and branching potential. (A)** Representative immunofluorescence image of day 4 definitive endoderm (DE) cells. Mean percentage of $SOX17^{+}$ nuclei (± SEM) from 3 independent experiments is indicated in green. Scale bar 100 µm. **(B)** Expression of selected marker genes on day 24 of differentiation normalized to GAPDH relative to day 0 (iPSCs). Bars represent mean fold change from 3 independent differentiation rounds. Error bars represent SEM. **(C)** Representative images of 3D branching organoids obtained by expanding lung epithelial progenitors in matrigel from day 24 of differentiation onwards without switching to a terminal differentiation medium. White arrow heads = bifurcation points. Scale bars 1000 µm (day 49, far left) and 400 µm (all others).

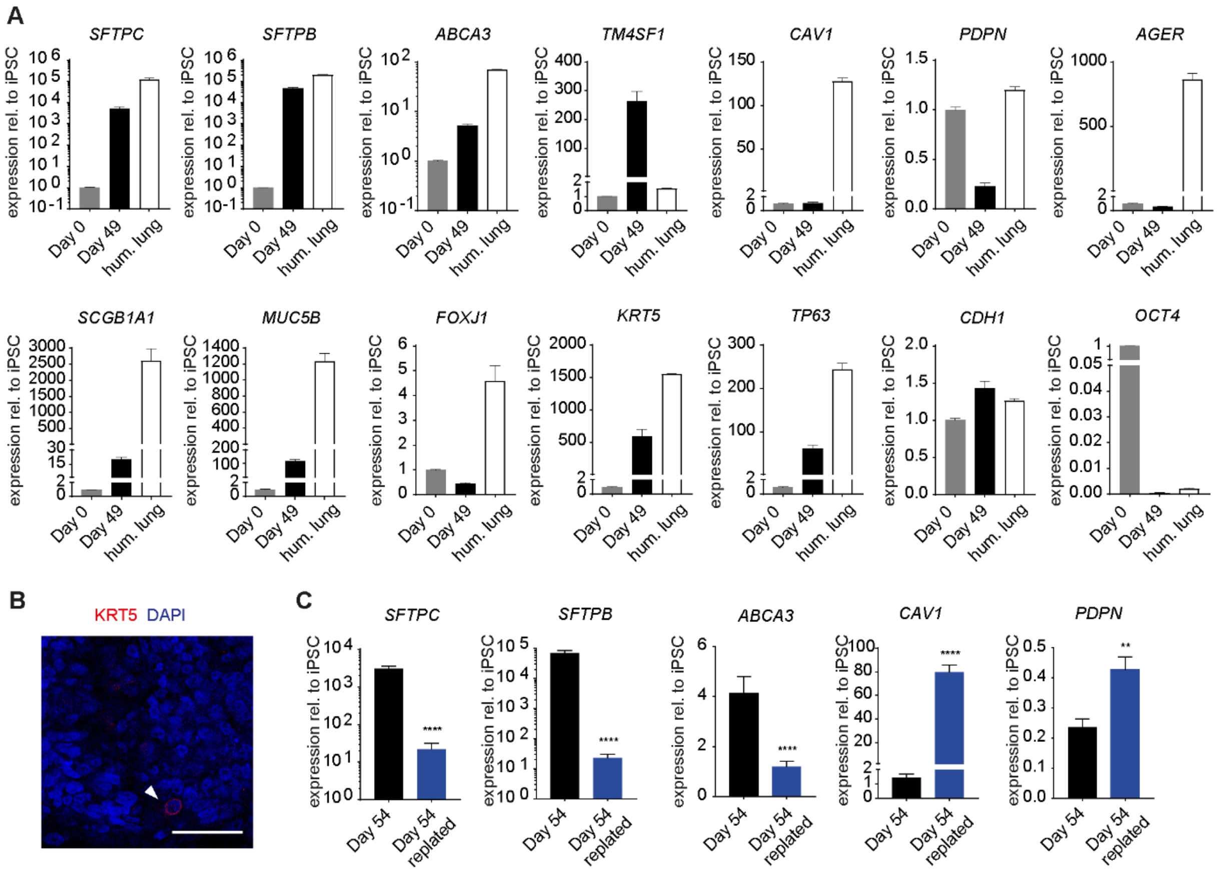


**Figure S2. Further characterization of iPSC derived ATII-like cells differentiated in air-liquid interface culture. (A)** Gene expression of alveolar and airway epithelial genes in iPSC derived ATII-like cells and in total human lung normalized to GAPDH relative to day 0 (iPSCs). Day 0 = iPSCs. Day 49 = iPSC derived ATII-like cells. Hum. lung = total human lung. Bars represent mean fold change from 9 independent differentiation rounds for iPSC derived cells. Error bars represent SEM. **(B)** Representative immunofluorescence against KRT5 in a day 49 culture (filled arrow head = rare $KRT5^+$ cell). Nuclei stained with DAPI (blue). Scale bars 50µm. **(C)** Gene expression of selected alveolar epithelial marker genes on day 54 of differentiation normalized to GAPDH relative to day 0 (iPSCs). Day 54 = iPSC derived ATII cells remained on transwell inserts at air-liquid interface culture. Day 54 replated = iPSC derived ATII cells replated onto tissue culture-treated plastic on day 49 and cultured under submerged conditions in 10% FBS medium. Bars represent mean fold change from 3 independent differentiation rounds. Error bars represent SEM. ** $P < 0.01$, **** $P < 0.0001$ by Mann-Whitney U-test.

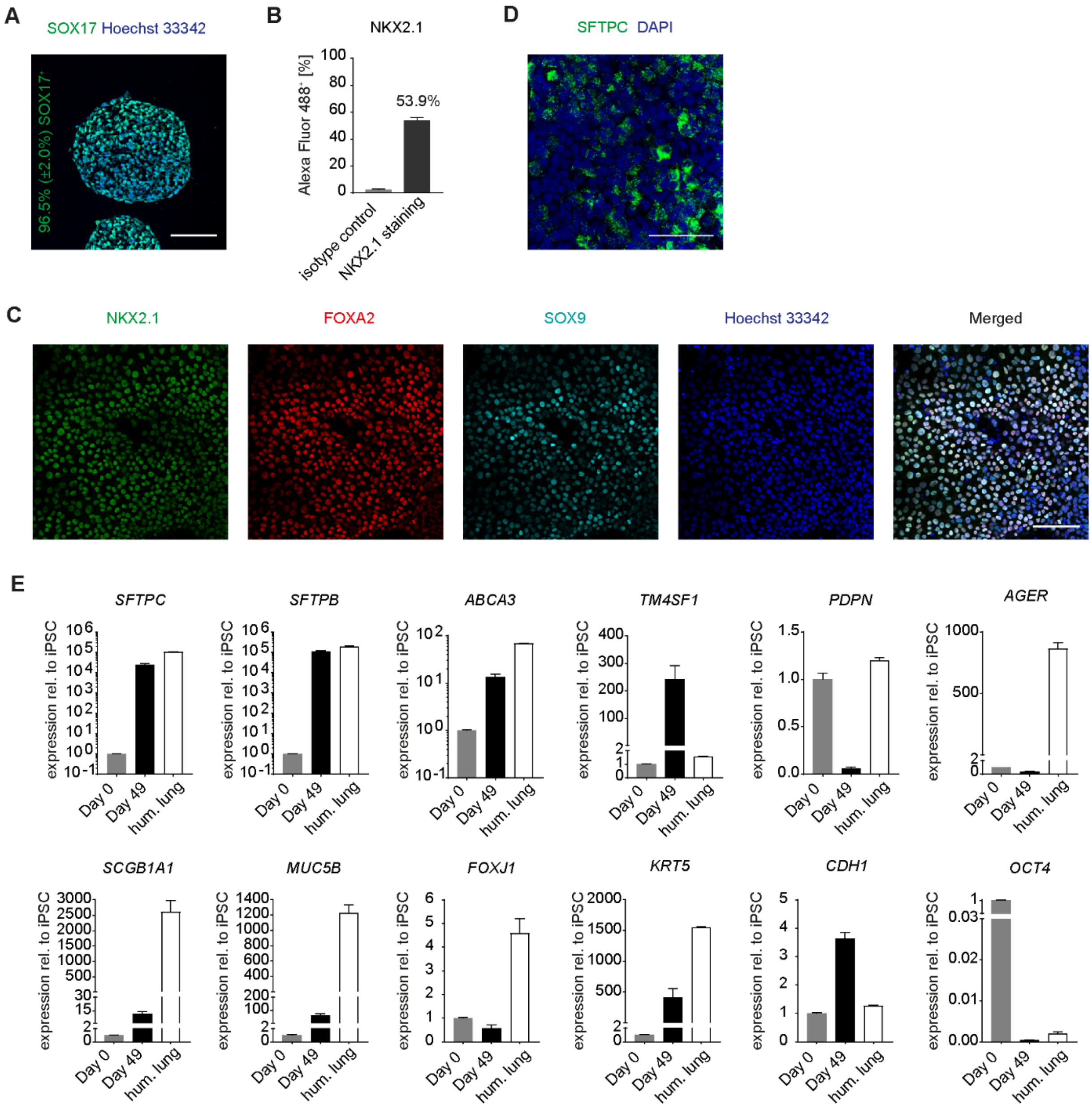


**Figure S3. Directed differentiation of an alternative iPSC line (SFC084-03-01) towards ATII-like cells. (A)** Representative immunofluorescence image of day 4 definitive endoderm (DE) cells. Mean percentage of SOX17$^+$ nuclei (± SEM) from 3 independent experiments is indicated in green. Scale bar 100 µm. **(B)** Day 24 representative flow cytometry analysis of NKX2.1$^+$ cells. Mean percentage of NKX2.1$^+$ cells from 3 independent differentiation rounds. Error bars represent SEM. **(C)** Representative immunofluorescence image of day 24 lung progenitors. Triple staining against NKX2.1 (green), FOXA2 (red) and SOX9 (cyan). Nuclei stained with Hoechst 33342 (blue). Scale bar 50µm. **(D)** Representative immunofluorescence image of a day 49 culture (whole insert). Staining against SFTPC. Nuclei stained with DAPI (blue). Scale bar 50µm. **(E)** Gene expression of alveolar and airway epithelial marker genes in iPSC derived ATII-like cells and in total human lung normalized to GAPDH relative to day 0 (iPSCs). Day 0 = iPSCs. Day 49 = iPSC derived ATII-like cells. Hum. lung = total human lung. Bars represent mean fold change from 3 independent differentiation rounds for iPSC derived cells. Error bars represent SEM.

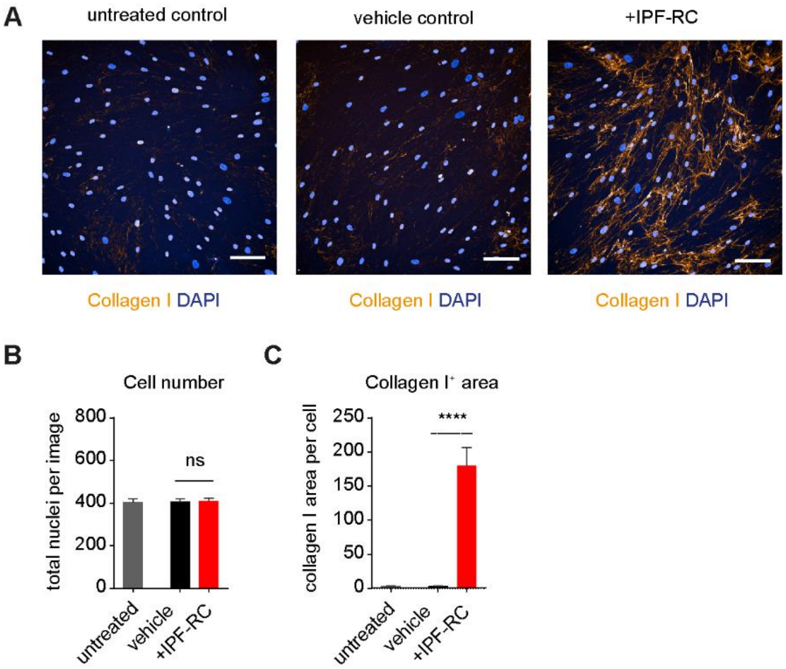


**Figure S4. IPF-RC induced collagen I formation in primary human lung fibroblasts. (A)** Representative immunofluorescence images against collagen I (orange) utilized for quantification. Nuclei stained with DAPI. Scale bars 100µm. Image based analysis of the number of nuclei **(B)** and collagen I **(C)** in primary human lung fibroblasts treated with IPF-RC for 72 h. Bars represent mean from 3 independent experiments. Error bars represent SEM. ns not significant, **** $P < 0.0001$ by Mann-Whitney U-test.

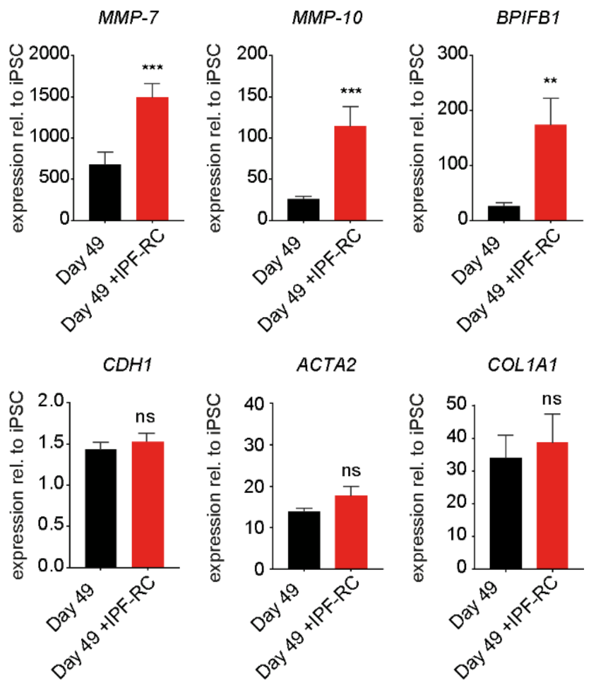


**Figure S5. The effect of IPF-RC treatment on gene expression of genes known to be deregulated in IPF including EMT-related genes in iPSC derived ATII-like cells.** qRT-PCR showing expression normalized to GAPDH relative to day 0 (iPSCs). Bars represent mean fold change from 6 independent differentiation rounds. Error bars represent SEM. ns not significant, * $P < 0.05$, ** $P < 0.01$ and *** $P < 0.001$ by two-tailed unpaired Student's t-test.

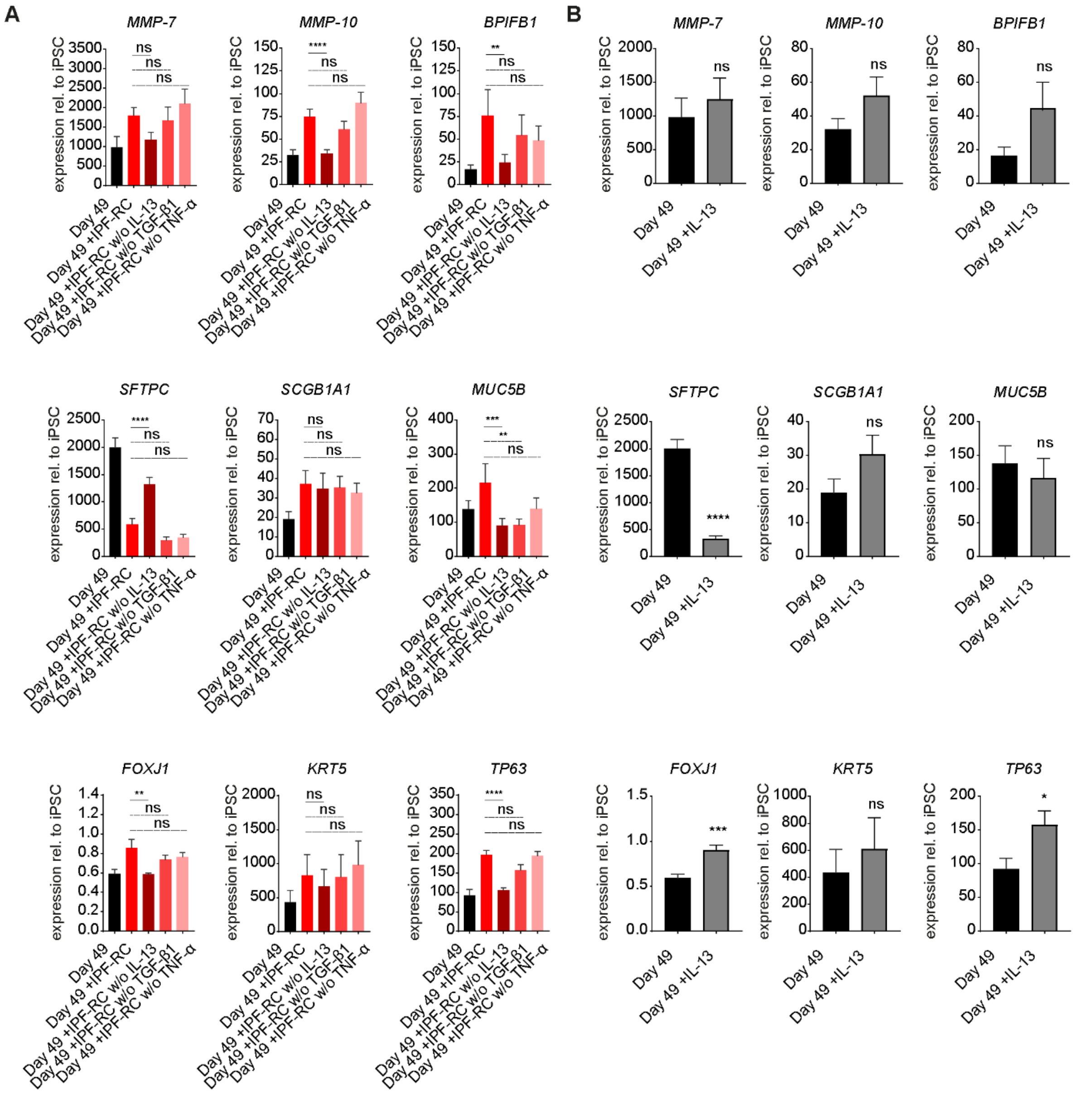


**Figure S6. Alterations in the effect of IPF-RC on iPSC derived ATII-like cell differentiation following removal of single cytokines from the cocktail and the effect of IL-13 alone.** Gene expression of IPF-related, alveolar and airway epithelial marker genes in iPSC derived ATII-like cells normalized to GAPDH relative to day 0 (iPSCs). Day 49 = iPSC derived ATII-like cells (control). **(A)** Removal of single cytokines from the cytokine cocktail. Day 49 +IPF-RC = iPSC derived cells treated with IPF-RC from day 35 of differentiation onwards. Day 49 +IPF-RC w/o IL-13 = iPSC derived cells treated with IPF-RC without addition of IL-13 from day 35 of differentiation onwards. Day 49 +IPF-RC w/o TGF-β1 = iPSC derived cells treated with IPF-RC without addition of TGF-β1 from day 35 of differentiation onwards. Day 49 +IPF-RC w/o TNF-α = iPSC derived cells treated with IPF-RC without addition of TNF-α from day 35 of differentiation onwards. Bars represent mean fold change from 3 independent differentiation rounds. Error bars represent SEM. ns not significant, ** $P < 0.01$, *** $P < 0.001$ and **** $P < 0.0001$ by two-way ANOVA and Tukey test. **(B)** Stimulation with IL-13 only at the same concentration as included in IPF-RC. Day 49 +IL-13 = iPSC derived cells treated with 2.5 ng/ mL IL-13 from day 35 of differentiation onwards. Bars represent mean fold change from 3 independent differentiation rounds. Error bars represent SEM. * $P < 0.05$, *** $P < 0.001$ and **** $P < 0.0001$ by Mann-Whitney U-test.

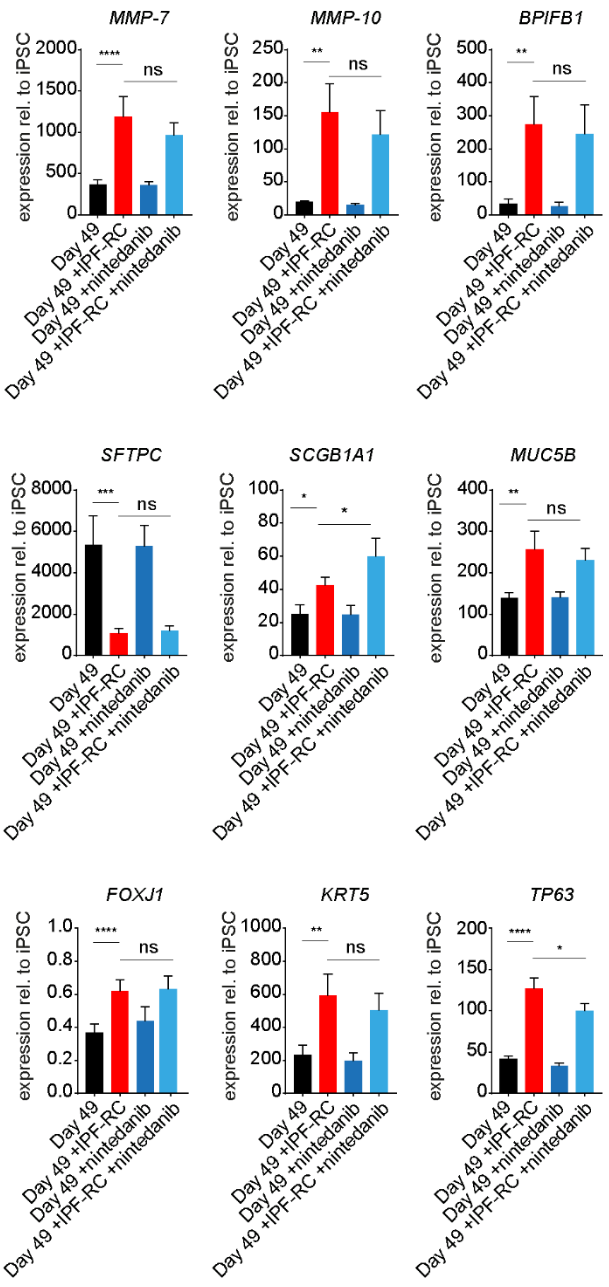


**Figure S7**. **The effect of nintedanib treatment on the transcriptional changes induced by IPF-RC treatment of iPSC derived ATII-like cells.** Gene expression of alveolar, airway epithelial and IPF-related marker genes in iPSC derived ATII-like cells and normalized to GAPDH relative to day 0 (iPSCs). Day 49 = iPSC derived ATII-like cells (control). Day 49 +IPF-RC = iPSC derived cells treated with IPF-RC from day 35 of differentiation onwards. Day 49 +100 nM nintedanib = iPSC derived cells treated with nintedanib from day 35 of differentiation onwards. Day 49 +IPF-RC +100 nM nintedanib = iPSC derived cells treated with IPF-RC and nintedanib from day 35 of differentiation onwards. Bars represent mean fold change from 3 independent differentiation rounds for iPSC derived cells. Error bars represent SEM. ns not significant, * $P < 0.05$, ** $P < 0.01$, *** $P < 0.001$ and **** $P < 0.0001$ by two-way ANOVA and Tukey test.

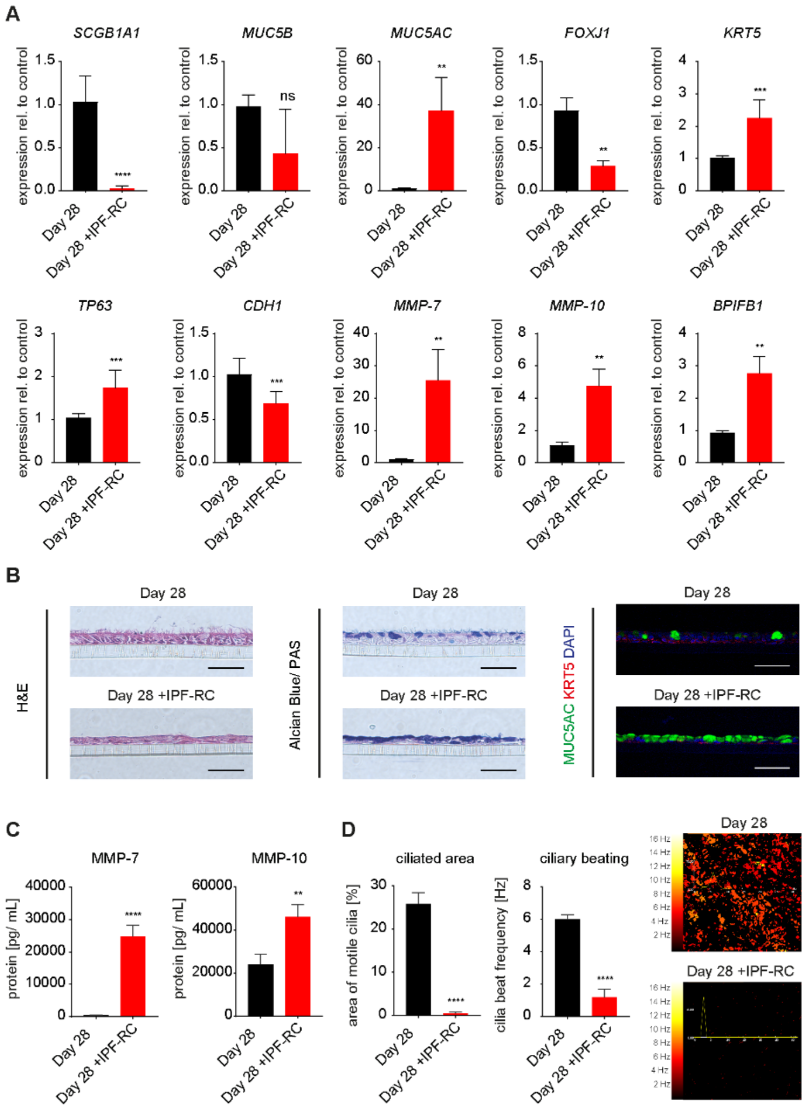


**Figure S8. IPF-relevant cytokine cocktail (IPF-RC) induced alterations in airway epithelial differentiation from primary human small airway basal cells. (A)** Gene expression in primary human small airway epithelial cell (SAEC) cultures normalized to GAPDH relative to untreated controls. Day 28 = SAECs differentiated for 28 days at air-liquid interface culture. Day 28 +IPF-RC = SAECs differentiated for 28 days at air-liquid interface in the presence of IPF-RC. Bars represent mean fold change from 3 independent experiments. Error bars represent SEM. ns not significant, ** $P < 0.01$, *** $P < 0.001$ and **** $P < 0.0001$ by Mann-Whitney U-test. **(B)** Representative haematoxylin and eosin stain (H&E, left), Alcian Blue/ PAS stain (middle) and immunofluorescence double stain (right) of representative cross-sections of primary human SAEC cultures. Immunofluorescence against MUC5AC (green) and KRT5 (red). Nuclei stained with DAPI (blue). Scale bars 50µm. **(C)** MMP-7 and MMP-10 protein levels in cell culture supernatant of SAEC cultures. Bars represent mean concentration from 3 independent experiments. Error bars represent SEM. ** $P < 0.01$, and **** $P < 0.0001$ by Mann-Whitney U-test. **(D)** Ciliary beat measurement in SAEC cultures. Left: Mean area covered by motile cilia and mean ciliary beat frequency calculated from 3 independent experiments. 6 regions per insert from 3 biological replicates were analyzed for each experiment. Error bars represent SEM. **** $P < 0.0001$ by Mann-Whitney U-test. Right: Representative regions analyzed for cilia beat measurement. Ciliary movement recording for 6 seconds per region.

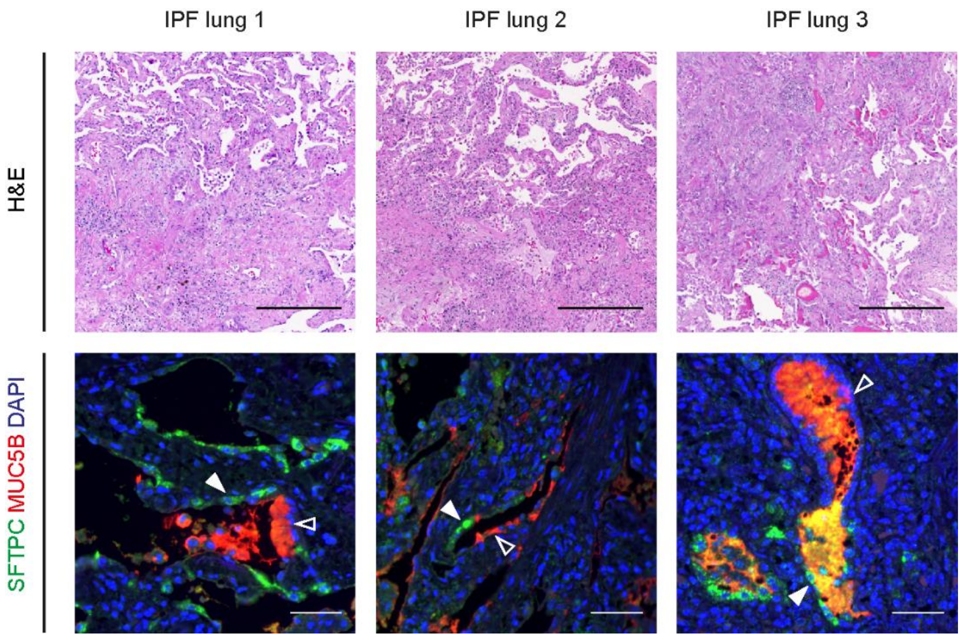


**Figure S9. Epithelial transition zones in cystic lesions of IPF patient lungs.** Representative haematoxylin and eosin stain (H&E) and immunofluorescence against SFTPC (green) and MUC5B (red) in three human IPF lungs. Epithelial transition zones in cystic lesions lined by $SFTPC^+$ cells (filled arrow heads) and $MUC5B^+$ cells (unfilled arrow heads). Nuclei stained with DAPI (blue). Scale bars 350µm in H&E images; 50µm in immunofluorescence images.

### SI TABLES

#### Table S1. Literature-based composition of an IPF-relevant cytokine cocktail (IPF-RC)

| Cytokine | Previous investigations of IPF BAL or sputum | IPF-RC [ng/mL] |
|---|---|---|
| TGF-β1 | • 32 pg/mL in IPF sputum (n=15), undetectable in healthy sputum (n=30) (14)<br>• 22.1 pg/mL in IPF with emphysema BAL (n=38); 14.4 pg/mL in IPF without emphysema BAL (n=64) (15)<br>• 1500 pg/mL in IPF BAL (n=16) (16) | 0.3 |
| IL-1β | • 0.6 pg/mL in IPF BAL (n=77), <0.3 pg/mL in healthy BAL (n=349) (17)<br>• 2.4 pg/mL in IPF BAL (n=11) (18) | 0.01 |
| TNF-α | • 1.9 pg/mL in IPF BAL (n=11) (18)<br>• 10 pg/mL in IPF sputum (n=15), 6.8 pg/mL in healthy sputum (n=30) (14)<br>• 0.87 pg/mL in IPF with emphysema BAL (n=38); 0.41 pg/mL in IPF without emphysema BAL (n=64) (15) | 0.1 |
| IL-8 | • 38.4 pg/mL in IPF BAL (n=11) (18)<br>• 276 pg/mL in IPF sputum (n=15), 46 pg/mL in healthy sputum (n=30) (14)<br>• 35 pg/mL in IPF BAL (n=13), 15 pg/mL in healthy BAL (n=9) (19)<br>• 179 pg/mL in IPF with emphysema BAL (n=38); 108 pg/mL in IPF without emphysema BAL (n=64) (15)<br>• 44.6 pg/mL in IPF BAL (n=7) (20) | 1.5 |
| MCP1 | • 70 pg/mL in IPF BAL (n=13), 5 pg/mL in healthy BAL (n=9) (19)<br>• 555 pg/mL in IPF with emphysema BAL (n=38); 351 pg/mL in IPF without emphysema BAL (n=64) (15) | 0.7 |
| IL-33 | • 4.13 pg/mL in IPF BAL (n=100), 1.15 pg/mL in healthy BAL (n=40) (21) | 0.04 |
| TSLP | • 2.5 pg/mL in IPF BAL (n=11) (18)<br>• 9.27 pg/mL in IPF BAL (n=100), 4.49 pg/mL in healthy BAL (n=40) (21) | 0.1 |
| IL-13 | • 250 pg/mL in IPF BAL (n=16), 60 pg/mL in healthy BAL (n=8) (22) | 2.5 |
| IL-4 | • 16 pg/mL in IPF BAL (n=16), 3 pg/mL in healthy BAL (n=8) (22) | 0.16 |

### Table S2. List of antibodies used in this study

| Antibody | Cat. No. | Vendor |
|---|---|---|
| ABCA3 (rabbit polyclonal) | ab99856 | abcam |
| Collagen type I (mouse IgG1) | SAB4200678 | Sigma-Aldrich |
| E-Cadherin (rabbit IgG) | 3195S | Cell Signaling |
| FOXA2 (goat IgG) | AF2400 | R&D systems |
| KRT5 (guinea pig polyclonal) | GP-CK5 | Progen |
| MUC5AC (mouse IgG1k) | MA1-38223 | Invitrogen |
| MUC5B (rabbit polyclonal) | HPA008246 | Sigma-Aldrich |
| NKX2.1/ TTF1 (mouse IgG1k) | MA5-13961 | Invitrogen |
| NKX2.1/ TTF1 (rabbit IgG) | WRAB-1231 | Seven Hills |
| Pro-SFTPC (rabbit polyclonal) | AB3786 | Millipore |
| SFTPB (rabbit polyclonal) | WRAB-48604 | Seven Hills |
| SFTPC (rabbit IgG) | HPA010928 | Sigma-Aldrich |
| SOX9 (rabbit IgG) | HPA001758 | Sigma-Aldrich |
| KRT5 (rabbit IgG) Alexa Fluor 647 conjugated | ab193895 | abcam |
| NKX2.1/ TTF1 (rabbit IgG) Alexa Fluor 488 conjugated | ab196470 | abcam |
| Donkey anti goat IgG Alexa Fluor 488 conjugated | A-11055 | Invitrogen |
| Donkey anti rabbit IgG Alexa Fluor 594 conjugated | A-21207 | Invitrogen |
| Donkey anti rat IgG Alexa Fluor 488 conjugated | A-21208 | Invitrogen |
| Goat anti guinea pig IgG Alexa Fluor 568 conjugated | A-11075 | Invitrogen |
| Goat anti mouse IgG Alexa Fluor 488 conjugated | A-11029 | Invitrogen |
| Goat anti mouse IgG1 Alexa Fluor 568 conjugated | A-21124 | Invitrogen |
| Goat anti mouse IgG Alexa Fluor 594 conjugated | A-11032 | Invitrogen |
| Goat anti rabbit IgG Alexa Fluor 488 conjugated | A-11034 | Invitrogen |
| Goat anti rabbit IgG Alexa Fluor 568 conjugated | A-11011 | Invitrogen |
| Goat anti rabbit IgG Alexa Fluor 647 conjugated | A-21245 | Invitrogen |
| Goat anti rat IgG Alexa Fluor 594 conjugated | A-11007 | Invitrogen |

**Table S3. List of TaqMan Gene Expression Assays used in this study**

| Target | Assay ID | Vendor |
|---|---|---|
| *ABCA3* | Hs00184543_m1 | Applied Biosystems |
| *ACTA2* | Hs00426835_g1 | Applied Biosystems |
| *AGER/ RAGE* | Hs00542584_g1 | Applied Biosystems |
| *AQP5* | Hs00387048_m1 | Applied Biosystems |
| *BPIFB1* | Hs00264197_m1 | Applied Biosystems |
| *CAV1* | Hs00971716_m1 | Applied Biosystems |
| *CDH1* | Hs01023895_m1 | Applied Biosystems |
| *COL1A1* | Hs00164004_m1 | Applied Biosystems |
| *FOXA2* | Hs00232764_m1 | Applied Biosystems |
| *FOXJ1* | Hs00230964_m1 | Applied Biosystems |
| *GAPDH* | Hs02758991_g1 | Applied Biosystems |
| *ID2* | Hs04187239_m1 | Applied Biosystems |
| *KRT5* | Hs00361185_m1 | Applied Biosystems |
| *MMP10* | Hs00233987_m1 | Applied Biosystems |
| *MMP7* | Hs01042796_m1 | Applied Biosystems |
| *MUC5AC* | Hs01365616_m1 | Applied Biosystems |
| *MUC5B* | Hs00861595_m1 | Applied Biosystems |
| *NKX2.1/ TTF1* | Hs00968940_m1 | Applied Biosystems |
| *PDPN* | Hs00366766_m1 | Applied Biosystems |
| *POU5F1* | Hs00999632_g1 | Applied Biosystems |
| *SCGB1A1* | Hs00171092_m1 | Applied Biosystems |
| *SFTPB* | Hs00167036_m1 | Applied Biosystems |
| *SFTPC* | Hs00161628_m1 | Applied Biosystems |
| *SOX17* | Hs00751752_s1 | Applied Biosystems |
| *SOX2* | Hs01053049_s1 | Applied Biosystems |
| *SOX9* | Hs00165814_m1 | Applied Biosystems |
| *TM4SF1* | Hs01547334_m1 | Applied Biosystems |
| *TP63* | Hs00978340_m1 | Applied Biosystems |

**Additional data table (separate file)**

Supplementary RNA-seq data

20. Vasakova M, Sterclova M, Kolesar L, Slavcev A, Pohunek P, Sulc J, Skibova J, Striz I. Bronchoalveolar lavage fluid cellular characteristics, functional parameters and

cytokine and chemokine levels in interstitial lung diseases. *Scandinavian journal of immunology* 2009; 69: 268-274.

21. Lee JU, Chang HS, Lee HJ, Jung CA, Bae DJ, Song HJ, Park JS, Uh ST, Kim YH, Seo KH, Park CS. Upregulation of interleukin-33 and thymic stromal lymphopoietin levels in the lungs of idiopathic pulmonary fibrosis. *BMC pulmonary medicine* 2017; 17: 39.

22. Park SW, Ahn MH, Jang HK, Jang AS, Kim DJ, Koh ES, Park JS, Uh ST, Kim YH, Park JS, Paik SH, Shin HK, Youm W, Park CS. Interleukin-13 and its receptors in idiopathic interstitial pneumonia: clinical implications for lung function. *Journal of Korean medical science* 2009; 24: 614-620.
